# A gene expression control technology for cell-free systems and synthetic cells via targeted gene silencing and transfection

**DOI:** 10.1101/2022.07.28.501919

**Authors:** Wakana Sato, Melanie Rasmussen, Nathaniel Gaut, Mahima Devarajan, Kaitlin Stokes, Christopher Deich, Aaron E. Engelhart, Katarzyna P. Adamala

## Abstract

Cell-free transcription-translation (TXTL) is an *in vitro* protein expression platform. In synthetic biology, TXTL is utilized for a variety of technologies, such as genetic circuit construction, metabolic pathway optimization, and building prototypes of synthetic cells. For all these purposes, the ability to precisely control gene expression is essential. Various strategies to control gene expression in TXTL have been developed; however, further advancements on gene-specific and straightforward regulation methods are still demanded. Here, we designed a novel method to control gene expression in TXTL, called a “silencing oligo.” The silencing oligo is a short oligonucleotide that binds to the target mRNA. We demonstrated that addition of the silencing oligo inhibits eGFP expression in TXTL in a sequence-dependent manner. We investigated one of the silencing oligo’s inhibitory mechanisms and confirmed that silencing is associated with RNase H activity in bacterial TXTL reactions. We also engineered a transfection system that can be used in synthetic cells. We screened two dozen different commercially available transfection reagents to identify the one that works most robustly in our system. Finally, we combined the silencing oligo with the transfection technology, demonstrating that we can control the gene expression by transfecting silencing oligo-containing liposomes into the synthetic cells.

## Introduction

Cell-free transcription-translation (TXTL) is a versatile protein expression platform used in synthetic biology. In response to a number of applications like genetic circuits^1,2^, metabolic engineering^3,4^, and communication networks^5^, there is an increasing demand for precisely controlled gene expression methods. Various technologies have been proposed to regulate protein expression and abundance in cell-free systems, such as transcription factors^2^, CRISPR technologies^6,7^, and the AAA+ family of proteases for tagged protein degradation^8,9^. However, to regulate specific genes towards constructing complex synthetic cells for biological research and practical applications, less complicated and quickly implemented methods without protein association need to be explored alongside current technologies.

Using antisense oligonucleotides to inhibit the mRNA translation has been intensively explored, especially in the research for therapeutic approaches^10–12^. Some antisense oligonucleotide drugs are currently on the market with FDA approval^13^. In the cellular environment, the mechanisms of antisense oligonucleotide driven inhibition include: (1) translational arrest, (2) RNase H1 associated mRNA degradation, (3) inhibition of 5’ cap formation, and (4) alternation of the splicing process^11,12^. Despite the tremendous number of studies in therapeutics, those antisense oligonucleotides’ functions in cell-free systems, such as *Escherichia coli* lysate-based TXTL, have not been clearly reported. Thus, we decided to characterize the inhibitory activity of different designs of antisense oligonucleotides and we aimed to develop a gene-specific inhibitor called a “silencing oligo” as a tool for cell-free systems and synthetic cells.

Transfection, using lipid reagents to facilitate the uptake of nucleic acids, is one of the most common ways to introduce foreign genes to live cells.^14,15^ In liposomal synthetic minimal cells, the only way to add genetic material after the formation of cells is to induce fusion with another liposome (which causes dilution of the cell content)^16,17^ or the use of cell-penetrating peptides carrying a nucleic acid payload (which has significant payload size limitations)^18^. Transfection enables the introduction of genetic material to induce or silence the expression of genes in tested cells. Engineering a transfection system for synthetic minimal cells is crucial to developing synthetic cells as a robust platform mimicking functions and processes of natural biology. Here we demonstrate a transfection system that works in one of the most commonly used synthetic cell liposome membranes. We demonstrate the use of this transfection system to deliver silencing oligos to regulate gene expression in synthetic cells.

## Results and Discussion

### Characterization of silencing oligos in cell-free systems

We first explored various oligo designs that showed sequence-specific inhibition in *Escherichia coli* (*E. coli*) TXTL.^19,20^ Our silencing oligos contain 14 or 15nt sequences complementary to their target mRNA, which is long enough to form a stable duplex at the TXTL reaction temperature of 30°C. We used eGFP to monitor the gene silencing activity because of the easiness of the measurement. We directly mixed silencing oligos into the TXTL reactions at a concentration of 10 μM, except in experiments comparing concentration dependency. We designed the silencing oligo to target different mRNA positions, such as the Shine-Dalgarno sequence, start codon, or chromophore coding region (**Supplementary Fig. 1**).

The design that showed the most potent inhibition of eGFP expression was “2OMe-oligo” (**Fig. 1a**). We replaced 2’-*O*-methylated RNA with DNA on both sides of the oligonucleotides (**Fig. 1abc**). The 2OMe-oligo showed a clear reduction in eGFP expression (**Fig. 1d**). We added this 2’-O-methylated RNA modification because the unmodified DNA oligonucleotides did not efficiently suppress eGFP (**Fig. 1d**, Unmod-oligos). We found some variation in the inhibition strength depending on what region was targeted. The strongest target was the chromophore targeting oligo with 0.04-fold eGFP expression compared to the control (**Fig. 1b**, 2OMe-Chrom). When we tested a longer oligo, 20nt, the inhibition activity increased and suppressed eGFP expression completely (**Fig. 1b**, 2OMe-Start+6bases). This increased inhibition is probably because of the oligo’s increased annealing temperature, which resulted in a more stabilized duplex formation in TXTL.

**Figure 1.**
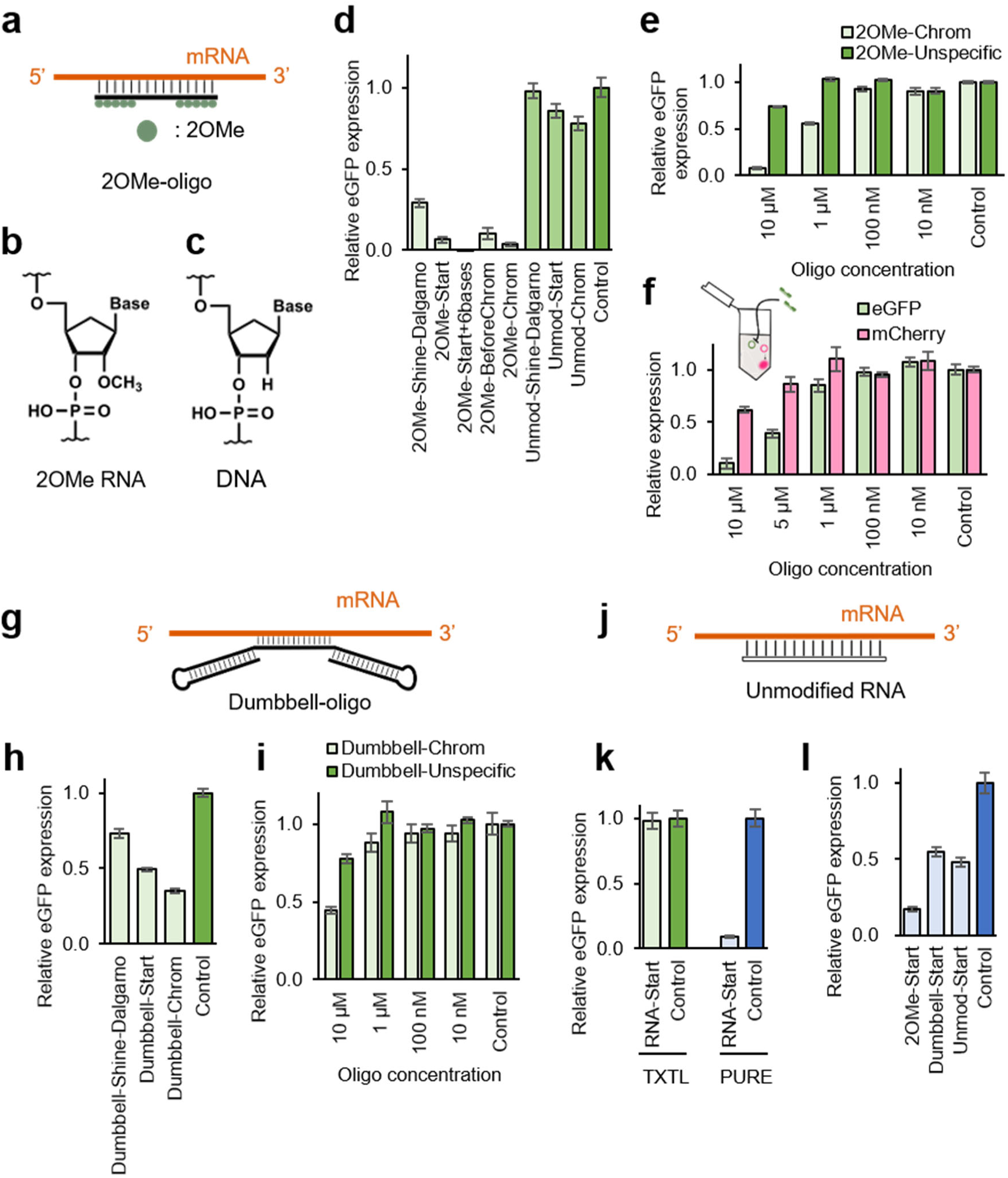
eGFP expression with different silencing oligos in cell-free systems. **a** The 2OMe-oligo consists of 2’-*O*-methylated RNA on both sides (green dots) and DNA in the middle of the structure. The oligo is 14 or 15nt length and contains a complementary sequence to the target mRNA. **b, c** Molecular structures of 2’-*O*-methylated RNA and DNA, respectably. **d** Inhibitory activity of 2OMe- and Unmodified DNA oligos in eGFP expression. **e** Sequence specificity of the 2OMe-oligo was tested with eGFP specific or unspecific 2OMe-oligo and eGFP fluorescence was measured. **f** Sequence specificity of 2OMe-oligo was tested with mCherry and eGFP plasmids in a single reaction. Only eGFP specific 2OMe-oligo (2OMe-Chrom) was added. The plasmid concentration was 10 nM for each, and the fluorescence was measured. **g** Dumbbell-oligo consists of DNA. Both sides of the oligo form duplex structures, and the middle sequence is designed to bind to the target mRNA. **h** Inhibitory activity of dumbbell-oligos in eGFP expression. **i** Sequence specificity of dumbbell-oligo was tested with eGFP mRNA specific or unspecific dumbbell-oligo and eGFP fluorescence was measured. **j** Unmodified RNA is 14nt ribonucleotides that contains the complement sequence of eGFP mRNA. **k** Comparison of eGFP expression in TXTL and PURE with 10 μM unmodified RNA oligo (RNA-Start). The inhibitory activity of the RNA oligo was only observed in the PURE reaction. **H** Silencing oligo’s inhibitory activity in eGFP expression in PURE. Silencing oligos were named in the order of “oligo structure – targeting position.” For the structure, the following abbreviations were used: “2OMe-”, 2’-*O*-methylated oligo; “Dumbbell-,” dumbbell oligo; “Unmod-,” unmodified DNA oligo; “RNA-,” unmodified RNA oligo. For the targeting sites, the following abbreviations were used: “Start,” start codon; “Start+6bases,” target start codon and 6 bases longer design (20nt); “BeforeChrom,” the sequence before the chromophore; “Chrom,” chromophore coding sequence; and “Unspecific,” random sequence. See the sequence map in **Supplementary Fig. 1** for the oligo targeting positions. All the silencing oligo sequences are in **Supplementary Table 1**. Throughout the graphs, “Control” represents eGFP expression in TXTL without a silencing oligo. Error bars indicate SEM, n=3. Original fluorescence data is in **Supplementary Fig. S25-S31**.

Then, we tested the sequence dependency of 2OMe-oligos to inhibit gene expression. We used the 2OMe-oligo containing an unrelated sequence to the eGFP mRNA. Although we observed a reduction in eGFP expression with unspecific 2OMe-oligo at 1 and 10 μM, sequence-specific inhibition was significant (**Fig. 1e**). We further tested the sequence dependency by expressing eGFP and mCherry in a single TXTL reaction. The 2OMe-oligo contains the eGFP mRNA’s complement sequence and does not bind with the mCherry mRNA. Although mCherry expression also decreased at oligo concentrations of 10 and 5 μM, the reduction of eGFP expression was more significant. These results demonstrated that 2OMe-oligos inhibit gene expression in TXTL sequence-dependently, with weaker off-target inhibition.

Another oligo design that inhibited eGFP expression in TXTL is called a “dumbbell-oligo” (**Fig. 1g**). We observed the strongest inhibition with the oligo targeting the chromophore resulting in 0.35-fold eGFP expression compared to the control (**Fig. 1h**, Dumbbell-Chrom). The dumbbell-oligos’ inhibition on eGFP expression was weaker than the 2OMe-oligo; however, they still showed significant inhibition and sequence-dependency when we compared eGFP specific and unspecific dumbbell-oligos (**Fig. 1i**).

We also tested some silencing oligos in the PURE system, another cell-free protein expression platform. PURE contains 36 purified enzymes involved in transcription and translation with purified 70S ribosomes^21^. Interestingly, when we tested silencing oligos in PURE, we observed that some designs worked better than in TXTL. The most prominent difference was from an unmodified RNA oligo that did not work in TXTL but only expressed 0.09-fold eGFP compared to the control (**Fig. 1jk**, **Supplementary Fig. 2** and **3**). The Unmodified DNA oligos also suppressed eGFP expression: 0.48-fold eGFP expression compared to the control, while those did not inhibit in TXTL (**Fig. 1dl**). The 2OMe- and dumbbell-oligos did not change the inhibition strength much (**Fig. 1dhl**).

Regarding the unmodified DNA oligo, we initially assumed the reason for non-inhibition was due to DNA degradation by nucleases in TXTL (**fig. 1d**). TXTL contains endogenous nucleases, and RecBCD is the major contributor to linear DNA fragment degradation in TXTL^22^. We tried two approaches to verify this assumption in two ways: adding RecBCD inhibitor GamS and using Akaby TXTL, a TXTL system without RecB. Both of them are verified to protect linear DNA in TXTL^19,22^. However, we did not see any enhancement of their inhibition activity on eGFP expression (**Supplementary Fig. 2** and **3**). Thus, the oligo degradation is not the reason for inefficient inhibition of unmodified DNA expression in TXTL. Potential explanations could be helicases or DNA binding proteins absent in the PURE reaction; however, more investigation is needed to get a clear explanation.

We also tested four other silencing oligo designs: a duplex-ends DNA oligo pair and two DNAzymes^23^ (**Supplementary Fig. 4**). Even though the other oligo-designs showed mild inhibition of eGFP expression in TXTL, it was not strong enough to declare that they can be used as inhibitors. The strongest design was the duplex-ends oligo targeting the chromophore with 0.49-fold eGFP expression compared to the control.

Additionally, we tried to see how plasmid and oligo ratios affect the inhibition of eGFP expression and tested varied ratios of eGFP plasmids and silencing oligos: 2OMe-oligo or dumbbell-oligo. At a plasmid concentration of 10 or 15 nM, the silencing activities of the 2OMe-oligo were consistent between the two concentrations (**Supplementary Fig. 5** and **6**). Thus, the silencing oligo concentration seems to determine the strength of the inhibitory activity rather than the ratio between the plasmids and oligos. The increase in variation and error is due to the lower concentrations of plasmids giving a lower eGFP fluorescence intensity. From the data of the plasmid concentration at 1 and 5 nM, we could not determine if a lower amount of template requires less oligo to inhibit expression.

### Investigation on the 2OMe-oligo working mechanism

Since the 2OMe-oligo showed the strongest gene silencing activity, we decided to explore its inhibition mechanism. First, we performed qPCR to measure mRNA abundance in the TXTL reactions with 2OMe-oligos. We performed qPCR with three different 2OMe-oligos: Pos.1, Pos.2, and Pos.3 (**Fig. 2a**). “Pos.1” binds at the start codon and the qPCR amplicon locates a region downstream of the oligo binding site; “Pos.2” binds in the middle and the qPCR amplicon contains the 2OMe-oligo binding site; “Pos.3” binds in the middle and the qPCR amplicon locates a region upstream of the oligo binding site.

**Figure 2.**
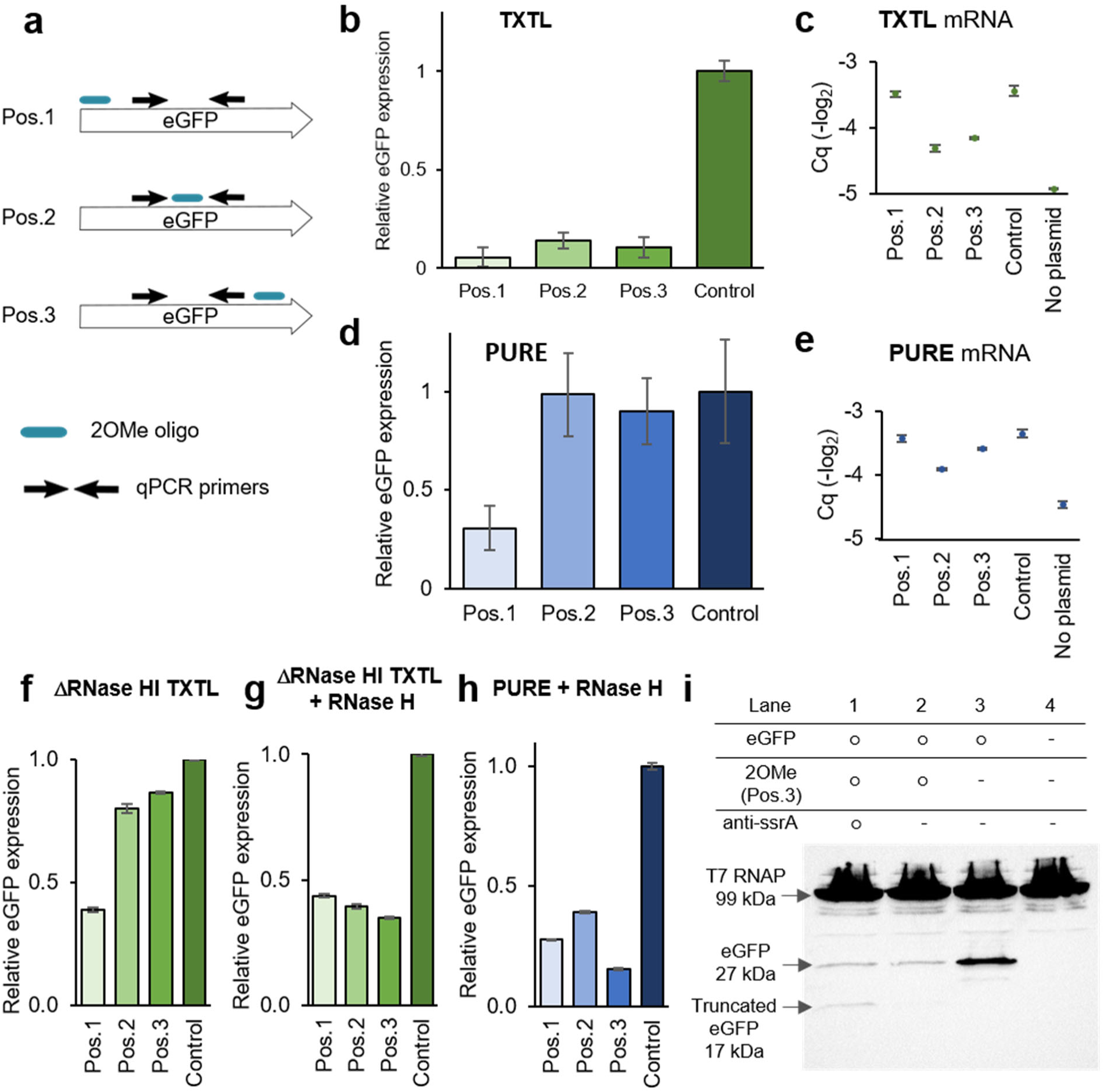
Investigations of 2OMe-oligos’ gene silencing mechanism. **a** The location relationships of the qPCR primers and silencing oligo target sites on mRNA. Light blue bars indicate the 2OMe-oligo binding sites (Pos.1, Pos.2, and Pos.3). Arrows indicate qPCR primer binding sites, which are consistent throughout Pos.1 to Pos.3. This figure only represents the locational relationships and does not correlate with the actual length of the eGFP gene. See **Supplementary Fig. 1** for the detailed binding locations on the gene. **b**, **d** eGFP expression with Pos.1-3 2OMe-oligos after 2 hours of TXTL (**b**) and PURE (**d**) reactions. **c**, **e** mRNA abundance with Pos.1-3 2OMe-oligo after 2 hours of TXTL (**c**) and PURE (**e**) reactions. **f** eGFP expression with Pos.1-3 2OMe-oligo in ΔRNase HI TXTL. **g**, **h** eGFP expression with Pos.1-3 2OMe-oligo in ΔRNase HI TXTL (g) and PURE (h). 0.05 units/μl RNase H (NEB, M0297L) was supplemented in the reactions. **i** Western blotting of TXTL reactions. TXTL reactions were fractionated on a 12% polyacrylamide gel for 90 minutes. N-terminal His-tagged T7 RNA polymerase is 99 kDa. Full-length N-terminal His-tagged eGFP is 27 kDa. The truncated eGFP by 2OMe-oligo interruption is 17 kDa. Anti-ssrA oligo is added to prevent tmRNA-associated truncated protein degradation. The table above the gel indicates which components are mixed in the TXTL reaction: “○” signifies the component was contained in the reaction and “ – ” signifies the component was not added. Control, eGFP expression in TXTL without silencing oligo; Cq, the quantification cycle. Error bars indicate SEM, n=3. Original fluorescence data is in **Supplementary Fig. S32-S33**. Original blot image is in **Supplementary Fig. S34**.

While we observed the 2OMe-oligos suppressed eGFP fluorescence in TXTL (**Fig. 2b**), mRNA abundance differed depending on the 2OMe-oligo targeting position (**Fig. 2c** and **Supplementary Fig. 7**). When using the Pos.1 oligo, the mRNA abundance was as high as the control-TXTL without 2OMe-oligo. However, the mRNA abundance significantly decreased when we used the Pos.2 or Pos.3 oligos. We also performed the same experiment in PURE. Surprisingly, when we used the Pos.2 or Pos.3 oligos, we did not see eGFP expression inhibition (**Fig. 2d**). The mRNA abundance with the Pos.1 or Pos.3 oligos was as high as the control, and the Pos.2 oligo slightly reduced the abundance (**Fig. 2e** and **Supplementary Fig. 8**).

The above results indicate that two mechanisms are involved in 2OMe-oligo silencing, dependent on where the 2OMe-oligo binds. A previous report also mentioned these mechanistic differences^24^. The inhibition mechanism may be irrelevant to the mRNA degradation for the Pos.1 oligo, which binds at the start codon. The mechanism is probably due to the steric hindrance of the 2OMe-oligo, which prevents ribosome translation initiation^10^. For the Pos.2 and Pos.3 oligos, which bind in the middle of the gene, the inhibition mechanism seems to be associated with mRNA degradation.

### RNase H involvement in 2OMe-oligo inhibition

RNase H1 is an endonuclease that cleaves RNA upon the hybridization with DNA. To see the effect of RNase H with the silencing oligos, we first added purified RNase H into the TXTL reaction (**Supplementary Fig. 9**). Adding RNase H in TXTL reduced eGFP expression efficiency (**Supplementary Fig. 9**, “None + RNase H” vs. “None”); however, we did not observe silencing oligo-dependent inhibition of expression. This observation initially surprised us because the widely used antisense oligo drugs require the RNase H activity to cleave and inactivate the gene expression of the targeted mRNA^11,12,25^.

We then hypothesized that endogenous RNase H in the extract already contributed to the silencing oligo inhibition. To confirm this, we removed RNase H from the cell extract. Although *E. coli* contains two RNase H genes (RNase HI and RNase HII, encoded by *rnhA* and *rnhB* genes, respectively), RNase HI is the only enzyme that attacks RNA:DNA hybrids *in vitro*.^26^ Thus, we proceeded to remove RNase HI from TXTL (ΔRNase HI TXTL). To make such a TXTL, we inserted an 8x-Histdine tag (His-tag) on the N-terminus of *rnhA* in the *E. coli* genome; we named the strain “Hina HI (<u>Hi</u>s-tagged R<u>Na</u>se <u>HI</u>)” (**Supplementary Fig. 10, Supplementary Fig. 11**). Using Hina HI allows us to remove His-tagged RNase HI during cell extract preparation by treatment with Ni-NTA beads.

In the ΔRNase HI TXTL, inhibition was reduced dramatically for the Pos.2 and Pos.3 oligos (**Fig. 2f**). This observation revoked what we saw in the PURE experiment (**Fig. 2d**). RNase HI may be the reason for the differences between TXTL and PURE (**Fig. 2b** and **2d**). To confirm this, we supplemented RNase H into the ΔRNase HI TXTL and PURE reactions and successfully restored the gene inhibitory activity of the Pos.2 and Pos.3 oligos (**Fig. 2gh**).

From these results, we think RNase HI cleaves mRNA upon binding of the silencing oligo, reducing gene expression. The mRNA abundances for the Pos.2 and Pos.3 reactions endorse this hypothesis: decreased mRNA in TXTL and high mRNA retention in PURE (**Fig. 2ce**). The potential slight decrease in mRNA abundance for the Pos.2 oligo in PURE might be because of the Pos.2 binding in the middle of the qPCR amplicon. That binding may prevent a PCR reaction and decrease the detectable amount of mRNA. 2OMe-oligo inhibitory activities in ΔRNase HI TXTL (**Fig. 2fg**) were lower than in the normal TXTL (**Fig. 2b**). This difference might come from batch differences or the ΔRNase HI procedure might affect TXTL composition. Since we saw a significant difference in the Pos.2 and Pos.3 oligo silencing activities with the addition of RNase H, we did not consider this a serious concern for understanding the RNase H involvement in the 2OMe-oligo gene silencing mechanism.

### tmRNA involvement in 2OMe-oligo inhibition

Considering the 2OMe-oligo mechanism, we wondered if we could detect truncated protein expression since the silencing oligo binds the middle of the gene. Ribosomes should translate the mRNA up to the 2OMe, even though eventually, the mRNA gets cleaved and degraded. To see if this translation occurs in TXTL, we used the Pos.3 oligo. The truncated protein length before the Pos.3 oligo binding site is 17 kDa, while the full-length N-terminus His-tagged eGFP is 27 kDa.

With the reaction containing the N-term His-tagged eGFP gene and the Pos.3 oligo, we did not see the expected size of the truncated product on a Western blot (**Fig. 2i**, lane 2). This suggests the truncated protein was either unproduced or degraded. When the mRNA is cleaved by RNases (including RNase HI), ribosomes get stalled in the middle of the mRNA. In such cases, *E. coli* can use the transfer-messenger RNA (tmRNA) system. ^27,28^ Briefly, tmRNA first binds to the stalled ribosome like transfer RNA and then adds a degradation tag (ssrA-tag) at the end of nascent synthesized truncated proteins. With the tmRNA system, ribosomes, mRNA, and truncated proteins can be released, and the incompletely synthesized proteins are degraded.

TXTL possesses the tmRNA system, and the supplementation of anti-ssrA oligonucleotides, which bind to the ssrA coding sequence of the tmRNA complex, can protect the synthesized truncated proteins from the tmRNA-associated degradations^29^. Therefore, we added anti-ssrA oligonucleotides into the reaction to protect the truncated proteins that were translated up to the 2OMe-oligo binding site. By adding anti-ssrA oligo, we could detect the truncated proteins in the TXTL reaction (**Fig. 2i**, lane 1). This result suggests that the tmRNA system associates with 2OMe-oligo gene silencing activity by degrading the truncated proteins and mRNA^27,28^. When we tried the anti-ssrA supplementation in ΔRNase HI TXTL, we could not observe the truncated proteins; thus, RNase HI cleavage seems required before the tmRNA-associated degradation (**Supplementary Fig. 12**).

### The 2OMe-oligo silencing mechanism in TXTL

Our observations suggested that at least two distinct mechanisms exist depending on the 2OMe-oligo binding position (**Supplementary Fig. 13**): (1) When the 2OMe-oligo binds near the start codon, the oligo prevents gene expression by preventing translation initiation^10^. (2) When the 2OMe-oligo binds the later sites downstream from the start codon, RNase H cleaves mRNA at the DNA/RNA hybrid position. Ribosomes translate the mRNA up to the mRNA binding site. The synthesized truncated proteins and the cleaved mRNAs are degraded through the tmRNA system and RNases (such as RNase E)^30^.

In an effort to identify RNases associated with 2OMe-oligo silencing, we also created ΔRNase III TXTL by using a strain “Hina III” (<u>Hi</u>s-tag R<u>Na</u>se <u>III</u>), which contains N-term His-tagged RNase III in its genome (**Supplementary Fig. 10**). RNase III is a double-stranded RNA-specific endonuclease. We chose to remove RNase III because the 2OMe-oligo consists of 2’-*O*-methylated RNA on both sides, and binding on mRNA results in RNA duplex formation that could be a substrate for RNase III. With ΔRNase III TXTL, we observed altered 2OMe-oligo silencing activity for the Pos.1 and Pos.3 oligos (**Supplementary Fig. 14**) and the truncated eGFP we saw upon anti-ssrA oligo supplementation disappeared (**Supplementary Fig. 15**). Thus, there is a possibility that RNase III also involves the 2OMe-oligo’s gene silencing activity; however, it can not be definitively determined from this experiment. This is because *rnc* (RNase III coding gene) and *era* (GTPase coding gene) are on the same operon and are two important genes for efficient ribosomal processing and maturation^31^. Affecting the operon by inserting a His-tag and a Kanamycin resistant gene (see method section) could significantly affect gene expression in *E. coli*, resulting in different TXTL compositions. Therefore, further investigation is required to determine the role of RNase III in the 2OMe-oligo silencing mechanism.

### Transfection in synthetic cells

To engineer a transfection system for the introduction of nucleic acids into lipid-based synthetic cells, we screened two dozen different commercially available transfection reagents (**Supplementary Table 2**). Different reagents are known to work best on specific cell lines and carry specific types of payloads. Many of the transfection reagents are known only under their brand names, with their specific lipid structures being trade secrets and not public knowledge. In this work, we obtained samples from most major brands of transfection reagents available in the US market, as well as samples of some less popular reagents reported to work well for challenging payloads.

We devised a screening pipeline to rapidly test for the transfection properties of different reagents. For those experiments, a sample of transfection reagent was mixed with the pUC19 plasmid, using the ratio and pre-incubation time recommended by the manufacturer instructions for each transfection reagent. The transfection sample was then mixed with POPC/cholesterol liposomes. After incubation, the liposomes were purified on a size exclusion column to remove unencapsulated DNA, the liposome membranes were lysed, and the samples were incubated with a restriction enzyme known to make a single cut on pUC19. After digestion, samples were analyzed on an agarose gel (**Supplementary Fig. 16-24**). This was a qualitative test, where the presence of digested DNA indicated that transfection was successful and pUC19 was delivered to the inside of liposomes. That test revealed a few transfection reagents that were capable of delivering a DNA plasmid payload into phospholipid vesicles under the tested conditions (**Supplementary Table 2**). Those transfection reagents were chosen for the next round of more stringent testing.

For the next round of transfection testing, we mixed transfection reagent with various payloads, both DNA and RNA: a circular pUC19, a linear pUC19 (linearized by a single cutting restriction digest), a smaller linear DNA fragment (500bp PCR product), a broccoli RNA aptamer, and a tRNA sample (*from E. coli* tRNA extract). In all cases, samples of the tested payload with the transfection reagent were incubated with liposomes and then samples were mixed with the Diamond Dye nucleic acid stain. The fluorescence of Diamond Dye significantly increases in the presence of oligonucleotides. We measured the fluorescence of the liposome transfection sample with Diamond Dye, then we lysed the liposomes and measured the sample fluorescence again. The measurement before lysis registered the fluorescence from Diamond Dye bound to nucleic acids outside of the liposomes, while the total fluorescence after the lysis measured the total amount of nucleic acids in the sample (oligos formerly inside and outside liposomes). Comparing those values, we can measure the amount of nucleic acids transfected inside the liposomes (**Fig. 3b-f**). The results indicated that all three transfection reagents tested in this round can deliver payloads inside lipid vesicles. We selected Lipofectamine 2000 for further study. None of the three tested reagents was significantly better at delivering all payloads, and Lipofectamine 2000 has a known chemical structure (3:1 mixture of DOSPA and DOPE) and is widely available across the world.

**Figure 3.**
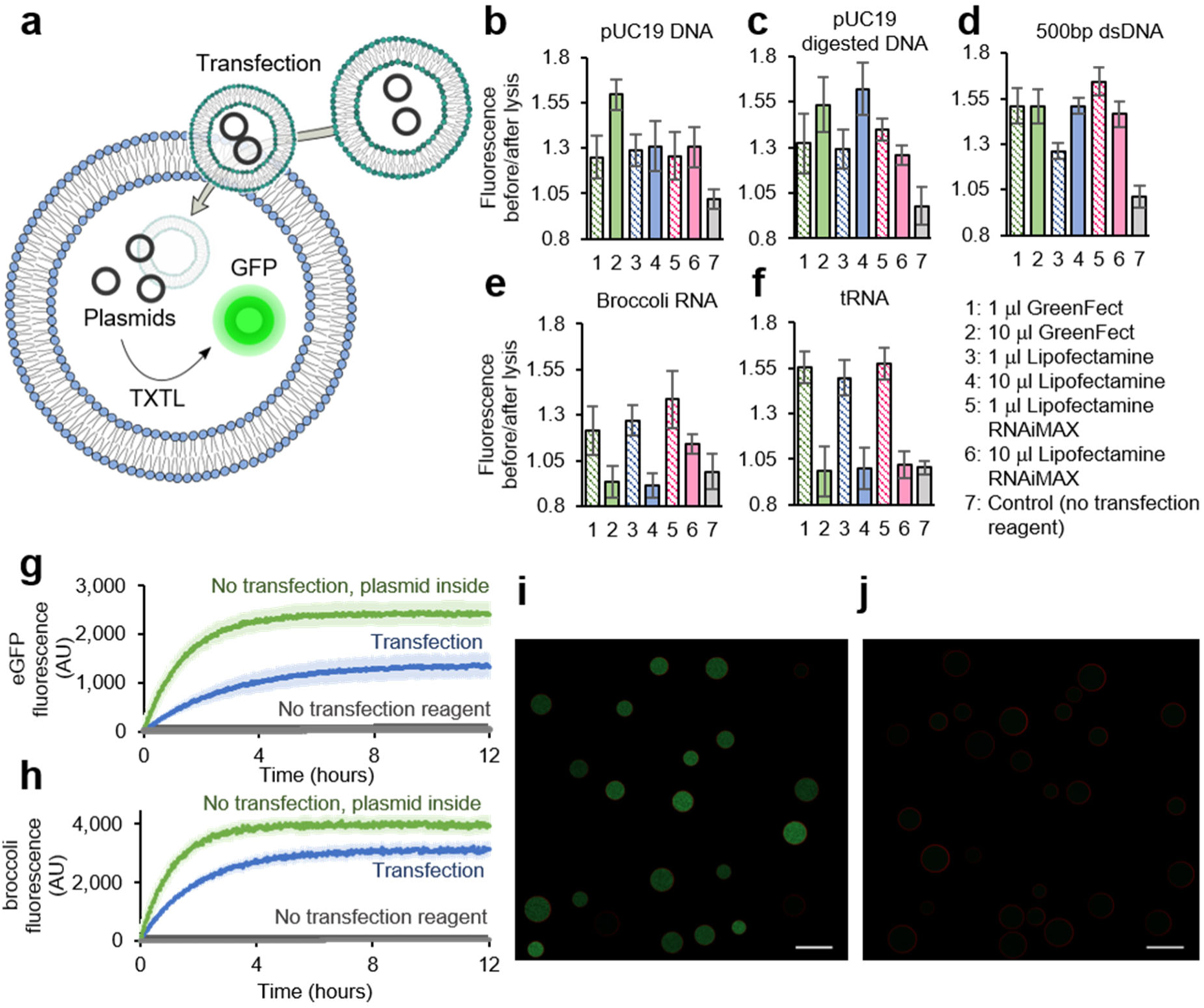
Transfection in synthetic cells. **a** General scheme of transfection in synthetic cell liposomes. Similar to natural cell experiments, the transfection reagent forms a complex with the nucleic acid payload and delivers that payload into the lumen of a phospholipid-based synthetic cell. Inside the synthetic cell, the transfected genes are expressed via a cell-free protein expression system encapsulated in the cell. **b-f** Delivery of different nucleic acid payloads to synthetic cell liposomes. GreenFect, Lipofectamine 2000 and Lipofectamine RNAiMAX reagents were tested in a Diamond Dye based assay. The fluorescence before lysis indicates fluorescence from nucleic acids outside liposomes and the fluorescence after lysis indicates fluorescence from all nucleic acids present in the sample; the ratio of those two values indicates the amount of nucleic acids that was present inside the liposome, i.e. the efficiency of transfection. The three transfection reagents were tested with **b** a circular plasmid, **c** a linear plasmid, **d** a 500bp dsDNA oligo, **e** a broccoli RNA aptamer, and **f** a tRNA mix. Error bars indicate SEM, n=3.Data points for each sample fluorescence before and after lysis are shown on **Supplementary Fig. 35** for DNA payloads and **Supplementary Fig. 36** for RNA payloads. **g** Time course for the transfection of broccoli aptamer DNA template inside synthetic cells capable of T7 RNA polymerase mediated transcription. The broccoli fluorescence was measured with an excitation wavelength of 472nm and emission wavelength of 507nm. Error bars indicate SEM, n=3. **h** Transfection of a DNA plasmid encoding a GFP gene into synthetic cells. The synthetic cells contain bacterial TXTL and T7 RNA polymerase. The GFP fluorescence was measured with an excitation wavelength of 472 nm and an emission wavelength of 507 nm. Error bars indicate SEM, n=3. Representative images of synthetic cells expressing GFP from transfected plasmid are on panel **i**, and a negative control without transfection reagents is on panel **j**. Scale bar is 10 μm. The synthetic cell membrane is dyed red with rhodamine dye.

We proceeded to use Lipofectamine 2000 to deliver DNA and RNA payloads inside synthetic cells. First, we prepared liposomes with the T7 RNA polymerase transcription system and we tested the delivery of a DNA template to produce a broccoli RNA aptamer (**Fig. 3g**). The broccoli fluorescence was recorded in liposomes, with a longer time to reach a steady-state than in the case of transcription from liposomes made containing broccoli template without the need for transfection (**Fig. 3g**). This delay is expected, as the transfection reagent needs to deliver the DNA template to liposomes before transfection can occur.

Next, we demonstrated that plasmid DNA can also be transfected. Synthetic cells containing bacterial TXTL cytoplasm were loaded with plasmids containing the eGFP gene (**Fig. 3h**). Similar to the transcription experiments, the GFP expression kinetics were slower in transfected samples than in samples that contained GFP plasmid from the beginning, consistent with the need to transfect before expression can begin. Microscope images confirm that only samples transfected with eGFP DNA express eGFP (**Fig. 3i**), compared to samples with eGFP DNA added to the outside of synthetic cells without the transfection reagents (**Fig. 3j**).

### Gene silencing in synthetic cells

Encouraged by the transfection experiments, we proceeded to use the newly established transfection system to deliver payloads of silencing oligonucleotides described earlier in this paper. We selected the two best performing silencing oligos tested on the eGFP gene, both targeting the eGFP chromophore: a 2OMe- and a dumbbell-oligo. We transfected each of those oligos into synthetic cells prepared with eGFP plasmids (**Fig. 4**).

**Figure 4.**
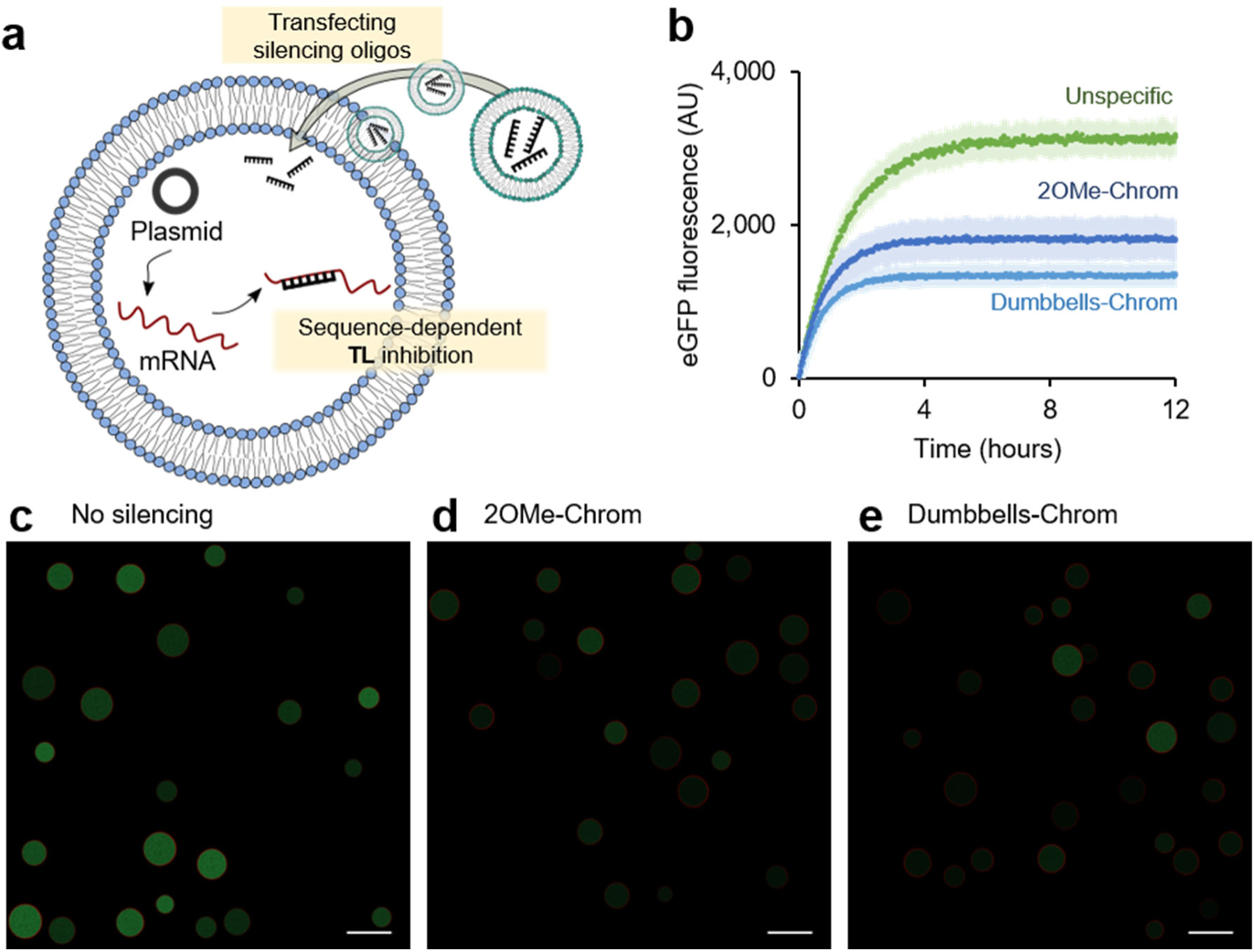
Gene silencing in synthetic cells. **a** Schematic image of transfecting silencing oligos to inhibit gene expression in a synthetic cell. **b** Time course of GFP expression in synthetic cells without silencing (transfected with short ssDNA not complementary to any part of the GFP gene), and transfected with either 2’OMe or with Stem-loop silencing oligo targeting GFP. The fluorescence was measured with an excitation wavelength of 472 nm and an emission wavelength of 507 nm. Error bars indicate SEM, n=3. Synthetic cells expressing GFP were transfected either with short ssDNA not complementary to any part of the GFP gene (panel **c**), with 2’OMe (panel **d**), or with the Stem-loop (panel **e**) silencing the oligo targeting GFP. The synthetic cell membrane is dyed red with rhodamine dye. Scale bar is 10 μm.

Both the 2OMe-oligo (**Fig. 4d**) and dumbbell-oligo (**Fig. 4e**) produced marked decreases in fluorescence in synthetic cells compared to synthetic cells transfected with a short DNA oligo not targeting eGFP (**Fig. 4c**). Quantification of the time courses of eGFP expression from those three synthetic cell samples showed that the initial kinetics for all three samples was similar, consistent with all the synthetic cell populations having the same amount of eGFP plasmid at the beginning of the experiment. As the transfection reagent delivered the silencing oligo, translation slowed, reaching steady states significantly lower than in the unsilenced sample (**Fig. 4b**).

### Summary and perspectives

The development of synthetic cells as a tool for biological research and practical applications requires a toolbox of methods for controlling gene expression past the point of formation of a synthetic cell. Here we demonstrated the regulation of gene expression via targeted silencing of genes in cell-free translation systems and in synthetic cells. We also demonstrated a method for introducing new genetic material into synthetic cells via transfection. Together, this work offers a robust toolbox for controlling gene expression in cell-free translation systems and in synthetic cells, providing a step towards engineering robust, life-like synthetic cells.

## Acknowledgments

This work was supported by: NASA award 80NSSC18K1139 Center for the Origin of Life - Translation, Evolution And Mutualism, John Templeton Foundation award 61184 Exploring the Informational Transitions Bridging Inorganic Chemistry and Minimal Life, NSF award 1844313 RoL: RAISE: DESYN-C3: Engineering multi-compartmentalised synthetic minimal cells, NSF award 2123465 Synthetic P-bodies: Coupling gene expression and ribonucleoprotein granules in synthetic cell vesicles for sensing and response, and Hackett Royalty Fund award. Wakana Sato was supported by the Funai Overseas Scholarship of The Funai Foundation for Information Technology

## Materials and Methods

### Materials

DNA oligonucleotides were purchased from Integrated DNA Technologies (IDT). Thermal cyclers used for sample incubation were Bio-Rad T100 thermocyclers running software version 1.201. The primer sequences are in **Supplementary Table 2**. eGFP and mCherry sequences are in **Supplementary Table 3**. C-terminal His-tagged eGFP plasmid was from our lab stock (Addgene, No. 178422)^32^. N-terminal His-tagged eGFP plasmid was from our lab stock (not deposited). mCherry plasmid was constructed under the T7Max promoter by using Epoch custom cloning service. pKD4 and pKM208 were purchased from Addgene^33,34^.

### Cell extract preparation

The cell extract preparation protocol was adapted from the Noireaux^19^ and Jewett^20^ protocols. The rosetta 2 cell extract preparation was followed by a method described previously^16^, with one modification. A 750 ml 2xYPTG culture was grown at 30°C instead of 37°C. Akaby cell extract preparation was described previously^22^. This process produces cell extract at a 3x concentration.

### TXTL reaction set up

Cell-free transcription-translation (TXTL) reactions were composed of the following: 12 mM Magnesium glutamate; 140 mM potassium glutamate; 1 mM DTT; 1.5 μM T7 RNA polymerase; 0.4 U/μl Murine RNase Inhibitor (NEB, M0314S); 1x cell extract; 1x energy mix; and 1x amino acid mix. The plasmid concentrations were 10 nM, and the silencing oligo concentration was 10 μM unless otherwise specified. The TXTL reactions were incubated at 30°C for 8 hours, followed by a 4°C hold until further procedures were performed.

10x Energy mix composition was the following: 500 mM HEPES, pH 8; 15 mM ATP; 15 mM GTP; 9 mM CTP; 9 mM UTP; 2 mg/mL E. coli tRNA; 0.68 mM Folinic Acid; 3.3 mM NAD; 2.6 mM Coenzyme-A; 15 mM Spermidine; 40 mM Sodium Oxalate; 7.5 mM cAMP; 300 mM 3-PGA.

10x amino acid mix was prepared by mixing 20 mM of the following amino acids: alanine, arginine, asparagine, aspartic acid, cysteine, glutamic acid, glutamine, glycine, histidine, isoleucine, leucine, lysine, methionine, phenylalanine, proline, serine, threonine, tryptophan, tyrosine, and valine. The amino acids were dissolved in pH 6.5, 400 mM potassium hydroxide solution.

### Hina HI and Hina III strains construction and cell extract preparation

An 8x His-tag was inserted on the N-term of *rnhA* (RNase H1 coding gene) or *rnc* (RNase III coding gene) on Rosetta 2 genome, together with kanamycin resistant gene (Km^R^) on the upstream of the target RNase (Supplementary Fig. 10).

We adapted a gene disruption method from a previous report^22^. Briefly, Km^R^ was amplified from pKD4 (Addgene, 45605) with primer 1 and primer 2 for Hina HI, and primer 3 and primer 4 for Hina III. The PCR products were transformed into pKM208 carrying Rosseta 2 (Millipore Sigma 71400-3) strain by electroporation. The successful mutants grew on LB agar plates with 50 μg/ml kanamycin. The mutation was verified by colony PCR with locus- and insert-specific primer pairs (see **Supplementary Fig. 10**). The successful mutants were stored as glycerol stocks.

The cell extract was prepared with a method used for the rosetta 2 cell extract preparation; however, the culture volume was 500 ml instead of 750 ml, and 50 μg/ml of kanamycin was used for pre-culture antibiotics instead of chloramphenicol. After all the procedures performed for the normal cell extract preparation, we treated the extract with Ni-NTA agarose beads (Goldbio, H-350-50) before aliquoting and freezing. 100 μl of Ni-NTA agarose beads was washed twice with 500 μl of water, then washed twice with wash buffer A (10 mM Tris-acetate pH 8.2, 14 mM magnesium acetate, 60 mM potassium acetate, 2 mM DTT). This wash buffer A was also used to suspend the cell pellet during cell extract preparation.^16^ The cell extract was mixed with the washed agarose beads and incubated for 1 hour on a horizontal rocker (Benchmark) at 4°C. After the incubation, the mixture of agarose beads and cell extract was loaded into spin-columns (Chrom Tech, CTF-CA020-01). This 3x cell extract was separated from the beads by spinning down with a tabletop centrifuge for 20 seconds. The resulting cell extract can be flash frozen and stored at -80°C.

### PURE reaction set up

We used PURExpress *In Vitro* Protein Synthesis Kit (NEB, E6800L) and the reaction was set up with the following manufacturer’s protocol. The plasmid concentration was 10 nM and the silencing oligo concentration was 10 μM. 1 U/μl of RNase inhibitor (NEB, M0314L) was supplemented in the reaction. After mixing all the components, the samples were incubated at 37°C for 8 hours followed by an end-point measurement.

### Fluorescence measurement

The eGFP fluorescence was measured at λ_ex_ 488 nm and λ_em_ 509 nm, and mCherry fluorescence was measured at λ_ex_ 587 nm and λ_em_ 610 nm, unless otherwise specified. The plate reader was set to a photomultiplier tube (PMT) “medium” and 6 reads per well. All fluorescent measurements were performed with SpectraMax. For the endpoint measurement of cell-free reactions, 19 μl of reaction was transported into a 384-well black flat bottom plate (Corning, 3575) after the incubation. For the kinetics measurement, 15 μl of cell-free reaction was incubated in the SpectraMax plate reader with a 384-well Black/ clear bottom plate (Corning, 4588), and the plate was sealed with a clear sealing tape to avoid evaporation.

### qPCR

The DNA in 2 μl of TXTL reaction was degraded with 0.5 μl of TURBO DNase (Invitrogen, AM2238) at 37°C for 30 minutes. The proteins in the TXTL were denatured by adding 15 mM EDTA and incubated at 75°C for 15 minutes. The denatured proteins were pelleted through centrifugation at 3,200 g for 2 min.

To prepare the 20 μl reverse transcription reaction, 2 μl of DNase-treated sample was mixed with 1 μM Primer 11, 10 mM DTT, 0.5 mM dNTP (Denville, CB4430-2), 5 U/μl protoscript II reverse transcriptase (NEB, M0368X), 1x protoscript II reverse transcriptase buffer, and 0.4 U/μl Murine RNase Inhibitor. The reverse transcription was performed at 42°C for 1 hour, followed by the inactivation at 65°C for 20 min.

The 25 μl qPCR reaction was performed by mixing the following: 1 μl of the reverse-transcribed DNA, 0.8 μM Primer 12, 0.8 μM Primer 13, 1x of OneTaq Hot Start 2X Master Mix with Standard Buffer (NEB, M0484L), and 1x Chai Green Dye (CHAI, R01200S). The qPCR was performed on CFX96 Touch Real-Time PCR Detection System (BioRad). The thermocycling program was the following: 1 cycle of 30 seconds denaturation at 95°C, 30 cycles of the following 15 seconds denaturation at 95°C, 15 seconds annealing at 50°C, 1 minute extension at 68°C, and 1 cycle of 5 minutes final extension at 68°C. The amplification curves plotted through CFX Maestro Software to determine Cq values and averages across 3 replicates of each sample were calculated separately.

### Western blot

TxTl reactions were fractionated on SDS-Page gel at 100V in 800ml 1x SDS running buffer (25mM Tris, 192mM Glycine, 3.5mM SDS). The gel percentage and fractionation time varied and are indicated on each figure. The gels were transferred to a 0.2 μ m nitrocellulose membrane at 100V for 1 hour in cooled transfer buffer (25mM Tris, 192mM Glycine, 10% methanol). The membrane was incubated with 5% nonfat milk in TBST for 1 hour followed by the addition of primary antibodies (BioLegend, 652505). After 1 hour of the primary antibody incubation, the membrane was rinsed three times with TBST followed by three 10 minutes washes in TBST. Secondary antibodies (BioLegend, 405306) were added to 5% nofat milk in TBST and incubate for 1 hour. After the incubation, the membrane was rinsed three times with TBST followed by three 10 minutes washes in TBST. The blots were imaged by ChemiDoc MP Imaging System with Image Lab Software (BIORAD), with the image application Blots, Chemi hi sensitivity reagent and Colorimetric. The chemiluminescent blot image and colorimetric image of the same blot were combined using the software merging function.

### Transfection experiments

All transfection reagents were mixed with DNA at the ratio described in the manufacturer’s instructions. All nucleic acid – transfection reagent complexes were pre-formed at room temperature for the amount of time described in the manufacturer’s instructions, then mixed with liposomes. Transfection reactions were incubated with gentle tumbling with incubation time of 4 hours for dye tests and 12 hours for gene expression tests. All liposomes were POPC/cholesterol unilamelar liposomes, prepared as previously described^16^.

### Relative expression calculation

To compare the silencing oligo inhibitory activity, we calculated the relative fluorescence protein expression as the following:

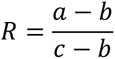

R: Relative expression

a: Sample fluorescence

b: Background fluorescence that is a cell-free reaction without plasmids and silencing oligos

c: None-oligo fluorescence that is a cell-free reaction with plasmids without silencing oligos

The errors of a ratio were calculated with the formula below:

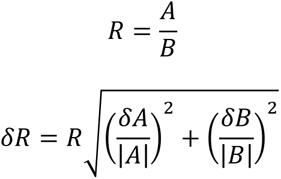

## Supporting materials

**Supplementary Figure 1.**
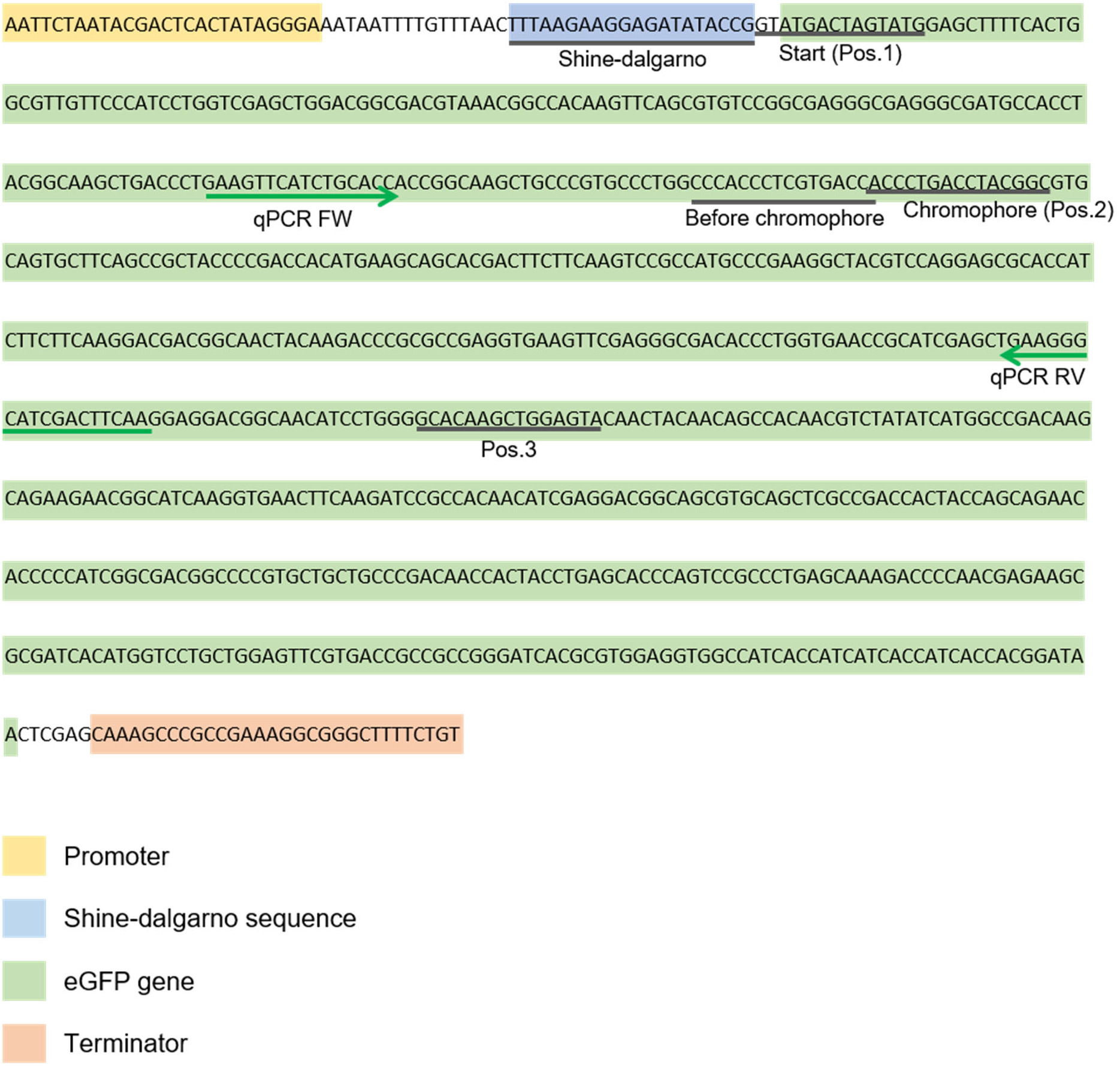
eGFP sequence map with silencing oligo binding positions. The part of DNA sequence from C-terminal His-tagged eGFP plasmid (Addgene, No. 178422). The silencing oligo binding positions are indicated with black lines with text labels. qPCR primers binding sequences are indicated with green arrows. The individual silencing oligo sequences are also listed in **Supplementary Table 1**.

**Supplementary Figure 2.**
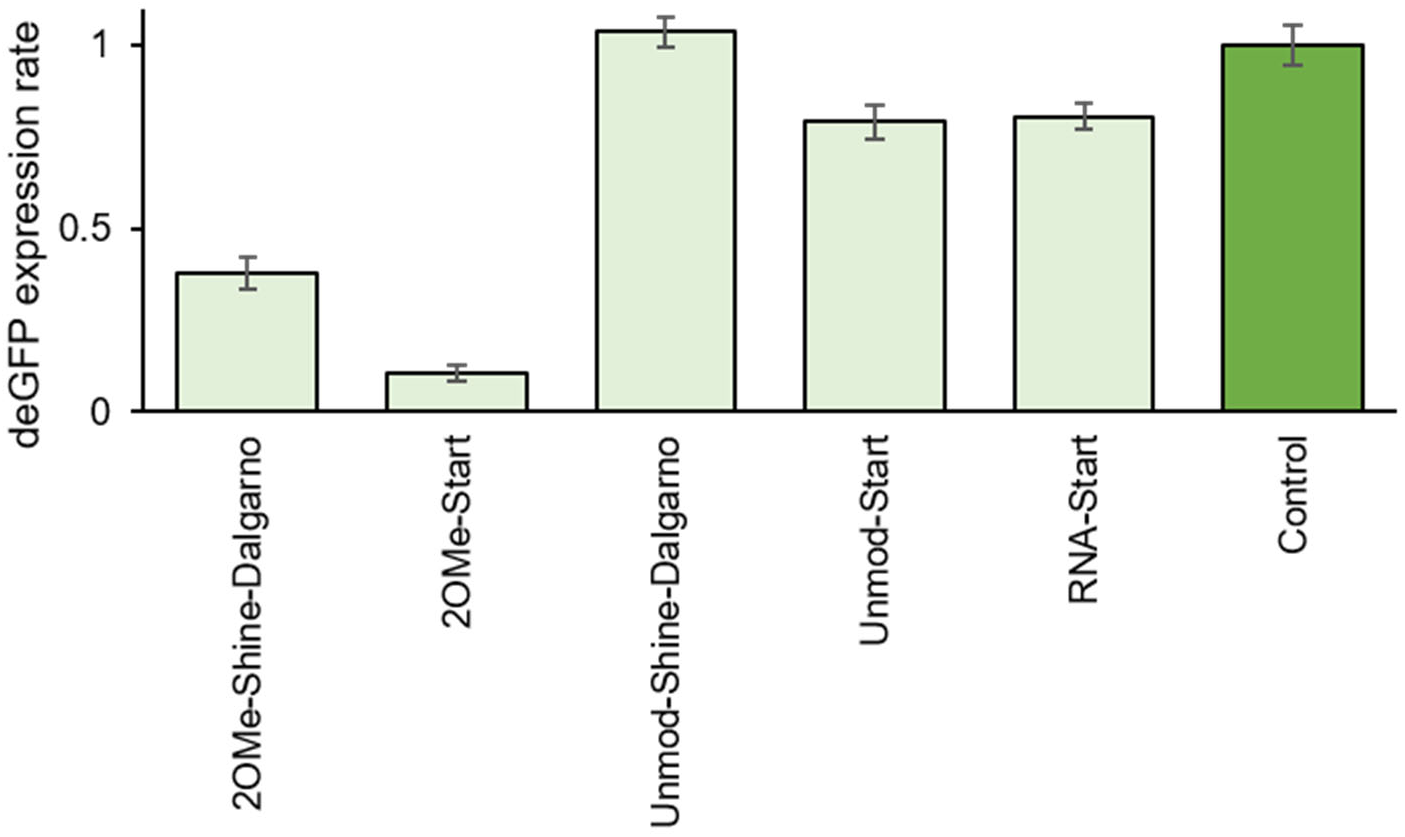
Silencing oligos’ eGFP inhibition activity in TXTL + GamS. eGFP was expressed with various silencing oligos in TXTL + GamS (3.5 μM). The silencing oligos were named in the order of “oligo structure – targeting position.” For the structure, the following abbreviations were used: “2OMe-,” 2-*O*-methylated oligo; “Unmod-,” unmodified DNA oligo; “RNA-,” unmodified RNA oligo. The start codon targeting was abbreviated as “Start.” Control reaction contains eGFP plasmid without silencing oligo. Error bars indicate SEM, n=3.

**Supplementary Figure 3.**
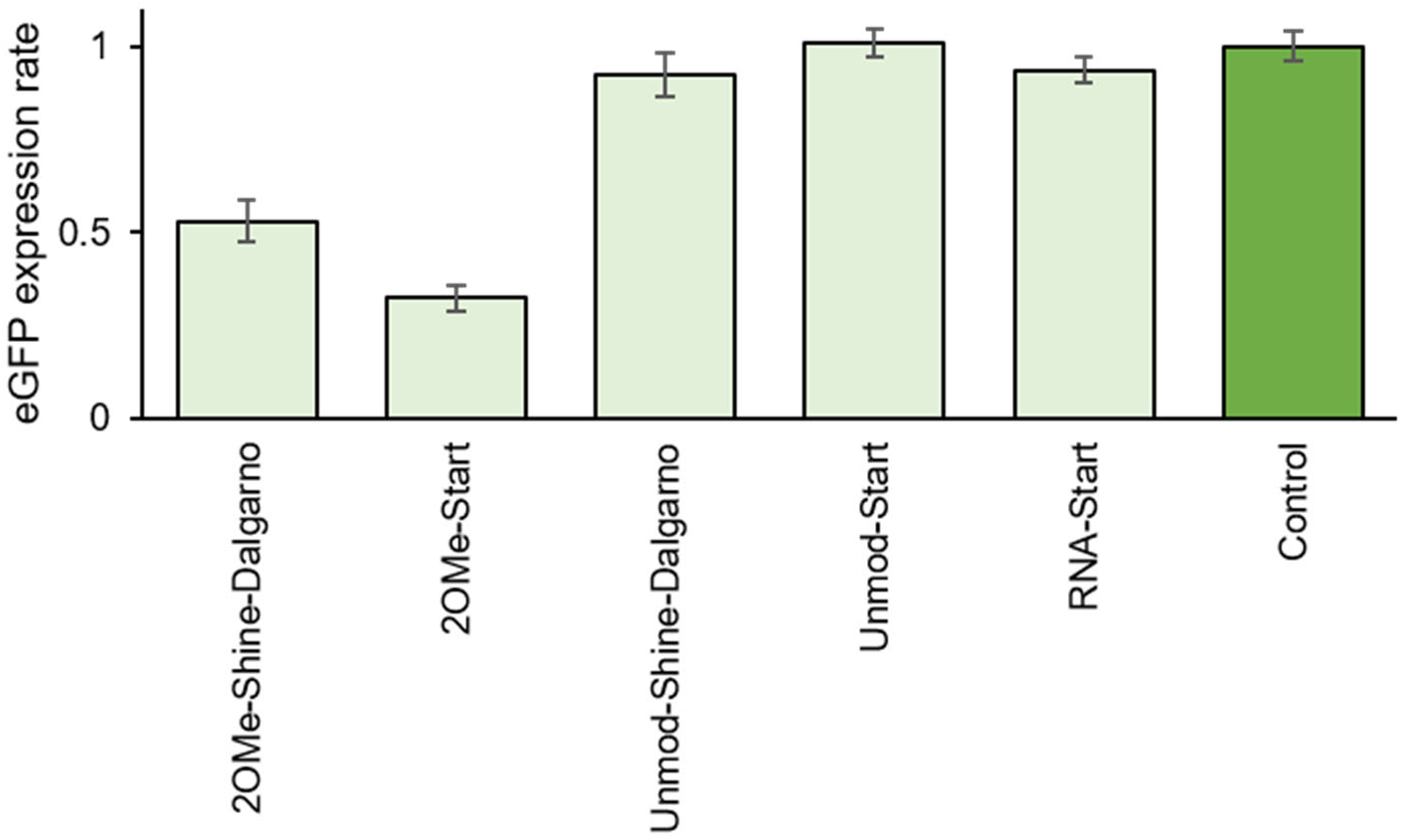
Silencing oligos’ eGFP inhibition activity in Akaby TXTL. eGFP was expressed with various silencing oligos in Akaby TXTL. Akaby is the Δ*RecB E. coli* strain. The silencing oligos were named in the order of “oligo structure – targeting position.” For the structure, the following abbreviations were used: “2OMe-”, 2-*O*-methylated oligo; “Unmod-,” unmodified DNA oligo; “RNA-,” unmodified RNA oligo. The start codon targeting was abbreviated as “Start.” Control reaction contains eGFP plasmid without silencing oligo. Error bars indicate SEM, n=3.

**Supplementary Figure 4.**
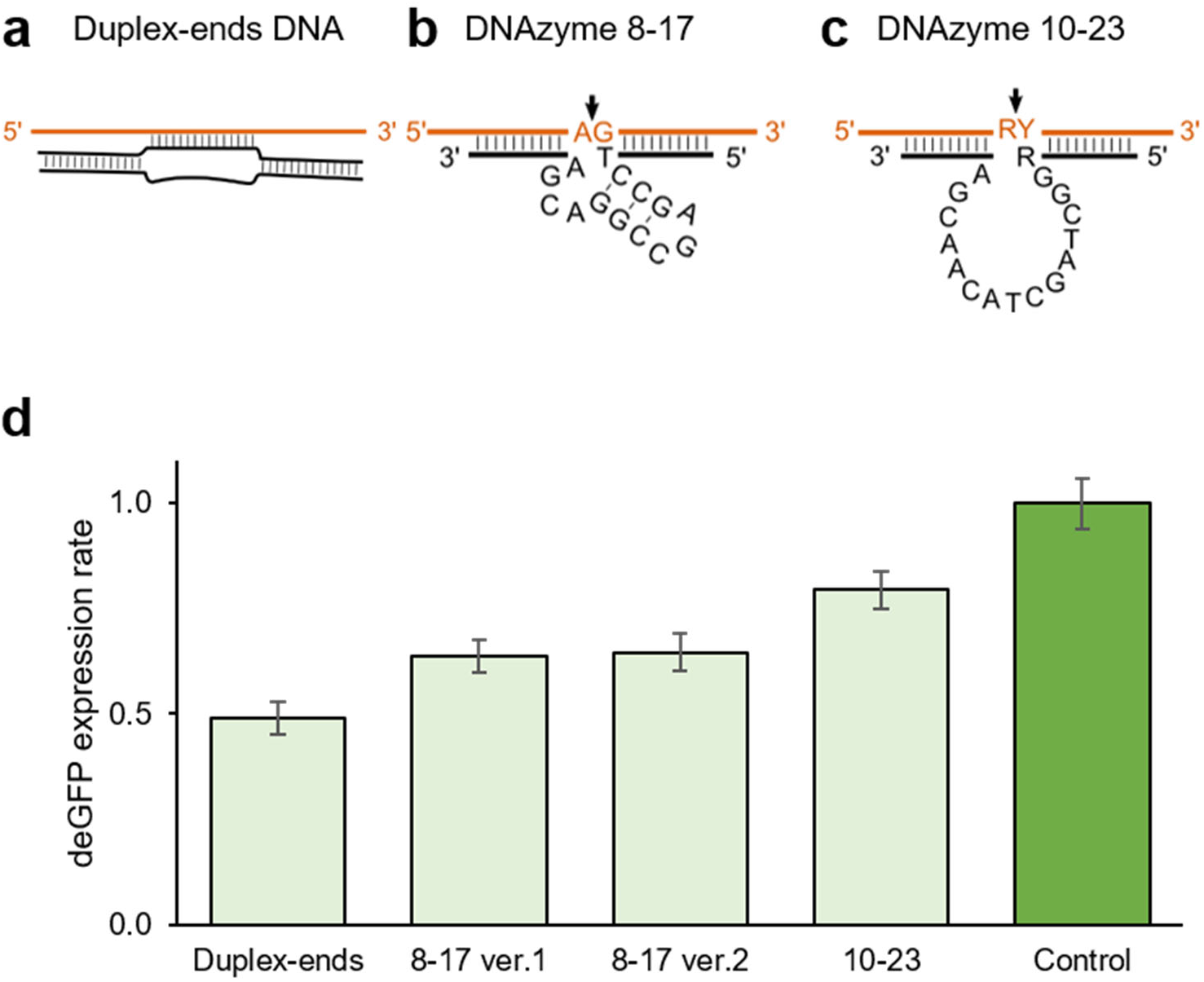
Other silencing oligo designs that tested. **a** Duplex-ends oligo consists of two DNA strands. One strand contains a complemental sequence to its target mRNA, and another contains a random sequence in the middle. Both edges are designed to form a duplex. **b**, **c** DNazyme design, the 8-17 catalytic motif (**b**) and the 10-23 catalytic motif (**c**). The figures were adapted from the Santoro and Joyce (1997) paper. The DNAzyme (bottom strand) binds to mRNA through Watson-Crick base pairing. Cleavage occurs at the position indicated by the arrow. R = A or G; Y = U or C. **d** eGFP was expressed with different designs of the silencing oligo in TXTL. For the 8-17 motif, ver.1 and ver.2 were designed to target different eGFP mRNA positions. Control reaction contains eGFP plasmid without silencing oligo. Oligo sequences are in **Supplementary Table 1**. Error bars indicate SEM, n=3.

**Supplementary Figure 5.**
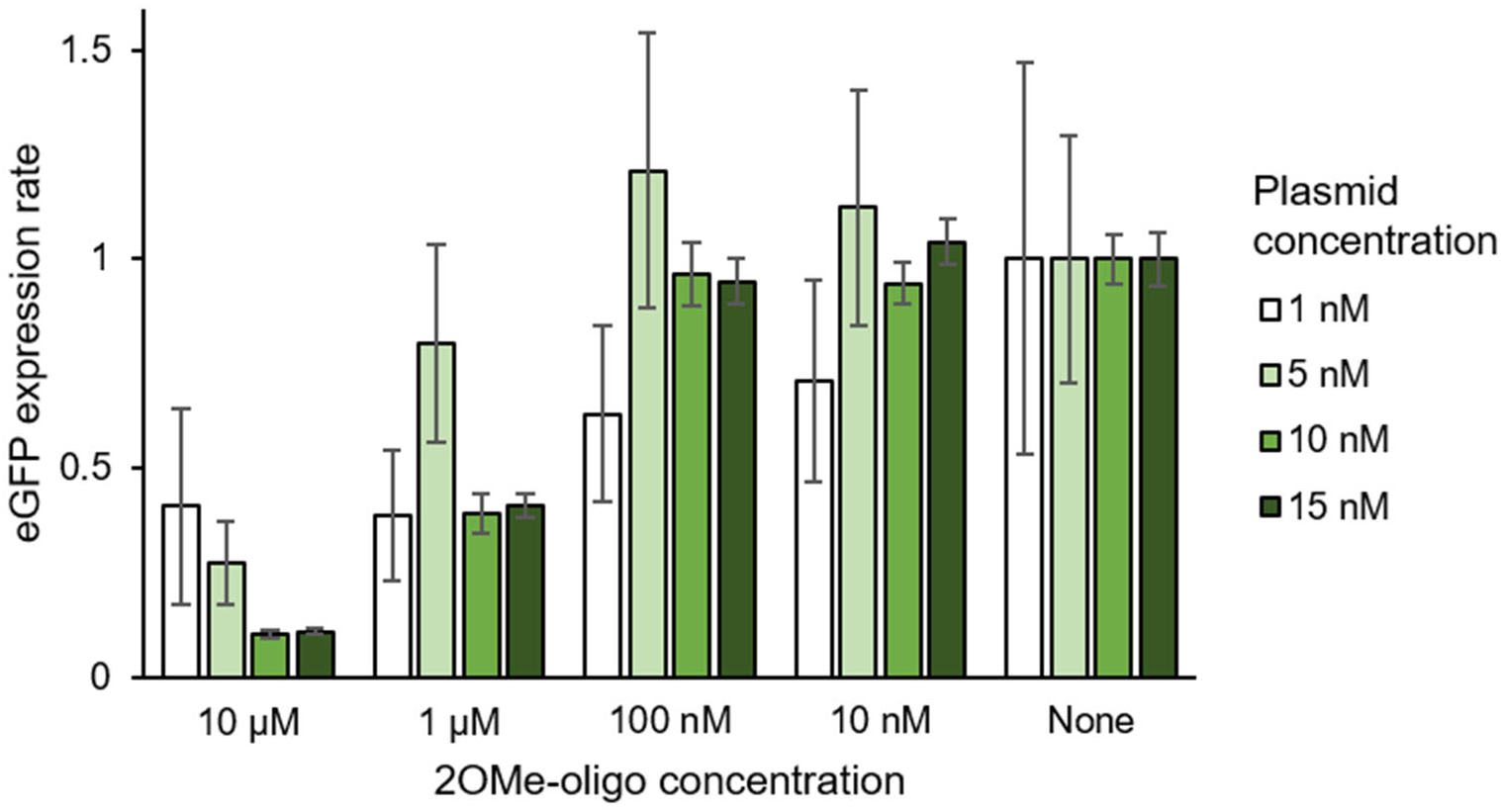
Various ratios of plasmid and oligo tested with 2OMe-oligo. eGFP expression with 2OMe-Chrom oligo. The eGFP plasmid concentrations were 1, 5, 10, and 15 nM. The eGFP expression efficiency was compared by varying oligo concentrations. Error bars indicate SEM, n=3.

**Supplementary Figure 6.**
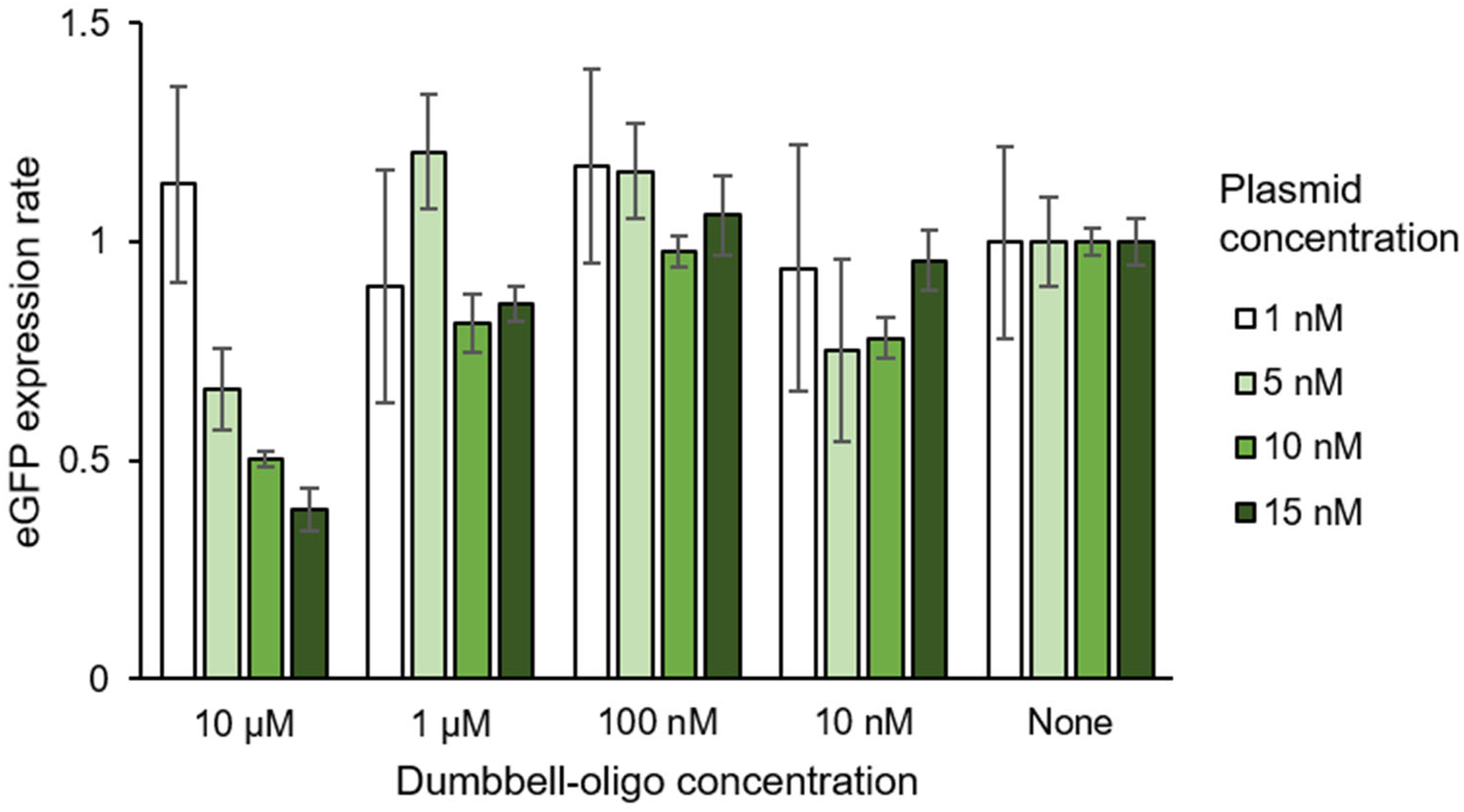
Various ratios of plasmid and oligo tested with dumbbell-oligo. eGFP expression with Dumbbell-Chrom oligo. The eGFP plasmid concentrations were 1, 5, 10, and 15 nM. The silencing oligo’s inhibition efficiency was compared. Error bars indicate SEM, n=3.

**Supplementary Figure 7.**
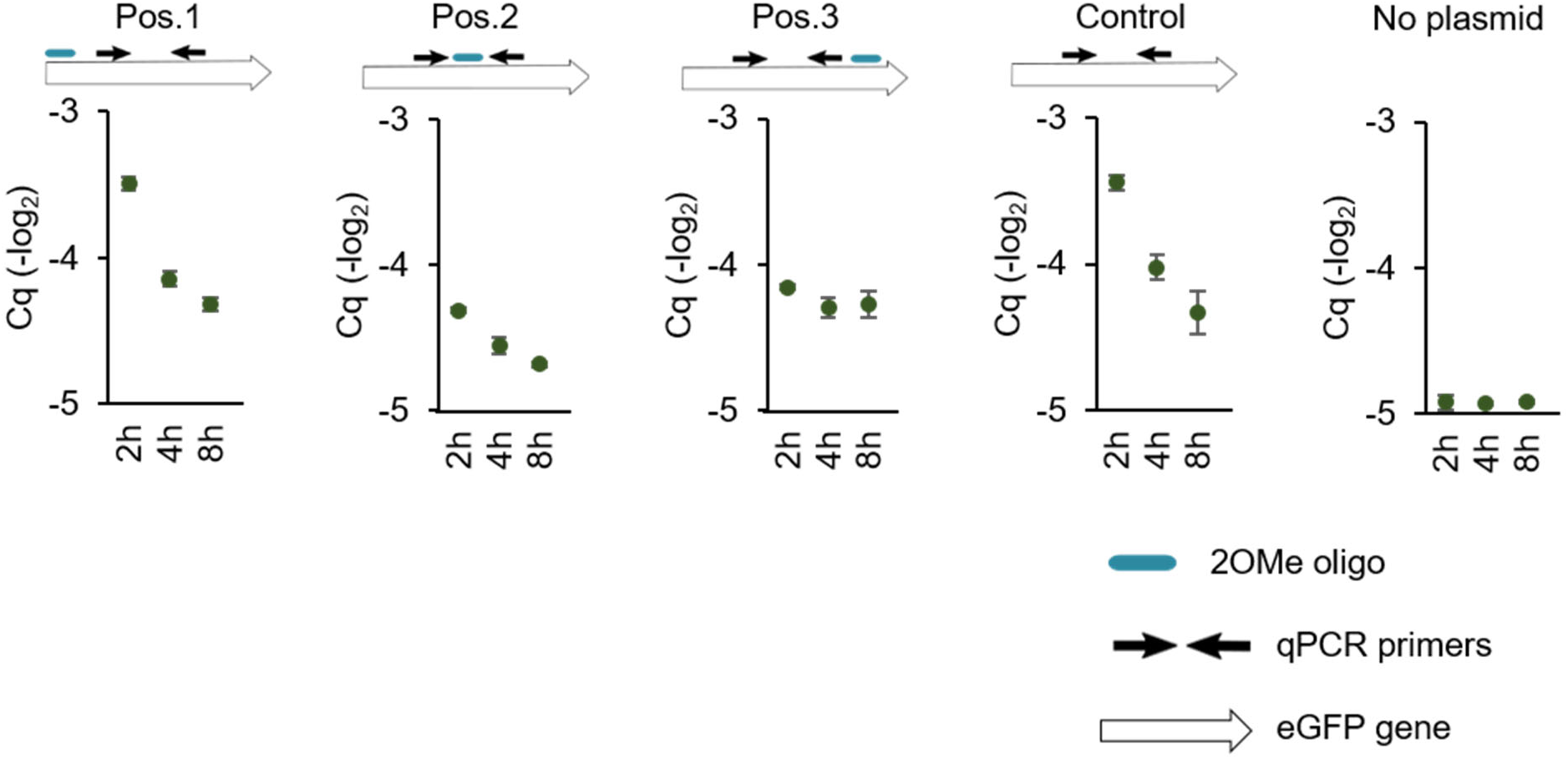
qPCR measurement of eGFP mRNA in TXTL, at 2, 4, and 8 hours. eGFP mRNA was measured in the TXTL reaction with 2OMe-oligos, Pos.1, Pos.2, and Pos.3. qPCR was performed after 2, 4, and 8 hours of incubation. The location relationships of the 2OMe-oligo binding position and qPCR amplicon are indicated above the graphs. Control is the reaction with eGFP plasmid without 2OMe-oligo. No plasmid is the reaction without eGFP plasmid. Error bars indicate SEM, n=3.

**Supplementary Figure 8.**
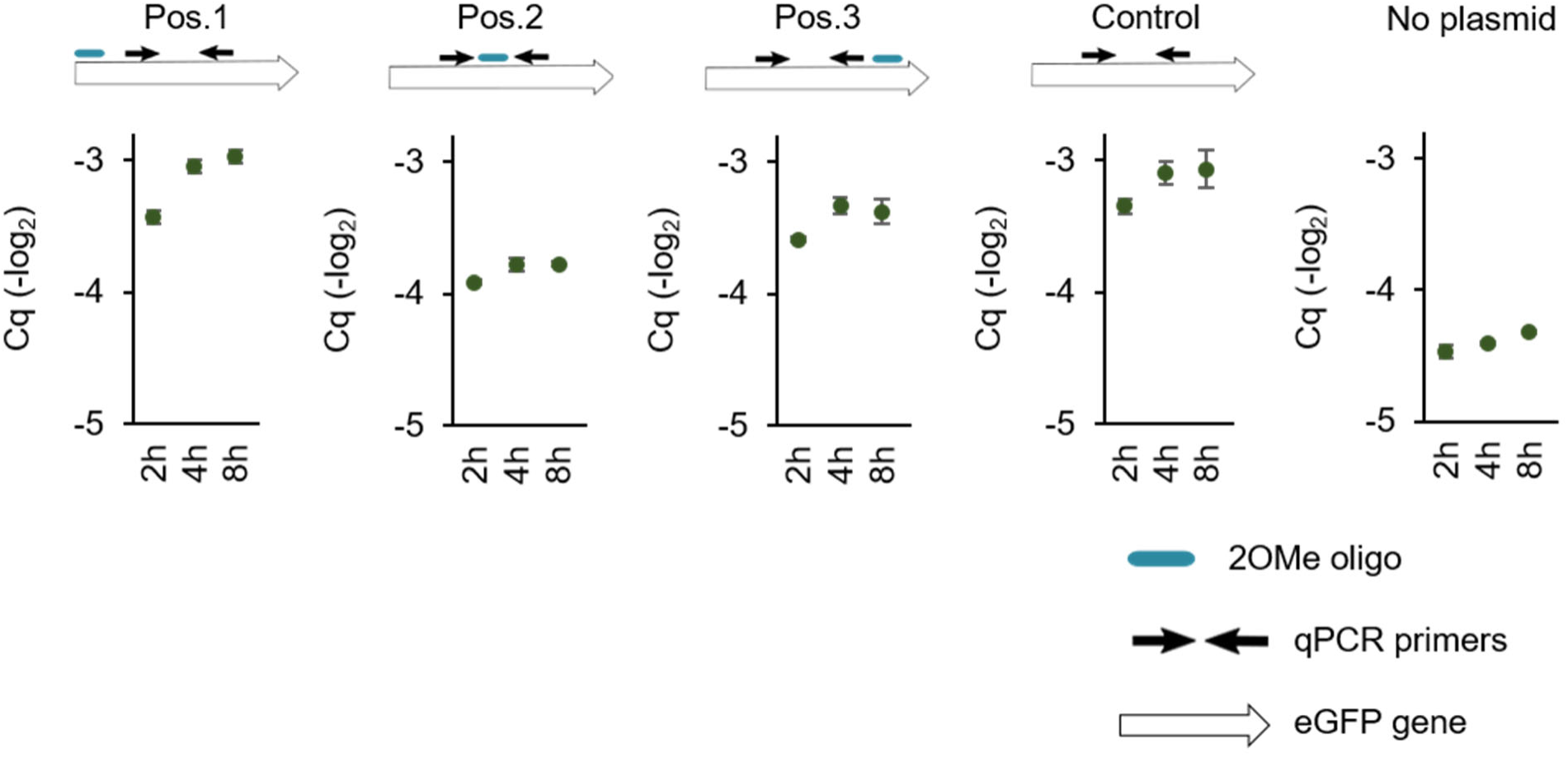
qPCR measurement of eGFP mRNA in PURE, at 2, 4, and 8 hours. eGFP mRNA was measured in the PURE reaction with 2OMe-oligo, Pos.1, Pos.2, and Pos.3. qPCR was performed after 2, 4, and 8 hours of incubation. The location relationships of the 2OMe-oligo binding position and qPCR amplicon are indicated above the graphs. Control is the reaction with eGFP plasmid without 2OMe-oligo. No plasmid is the reaction without eGFP plasmid. Error bars indicate SEM, n=3.

**Supplementary Figure 9.**
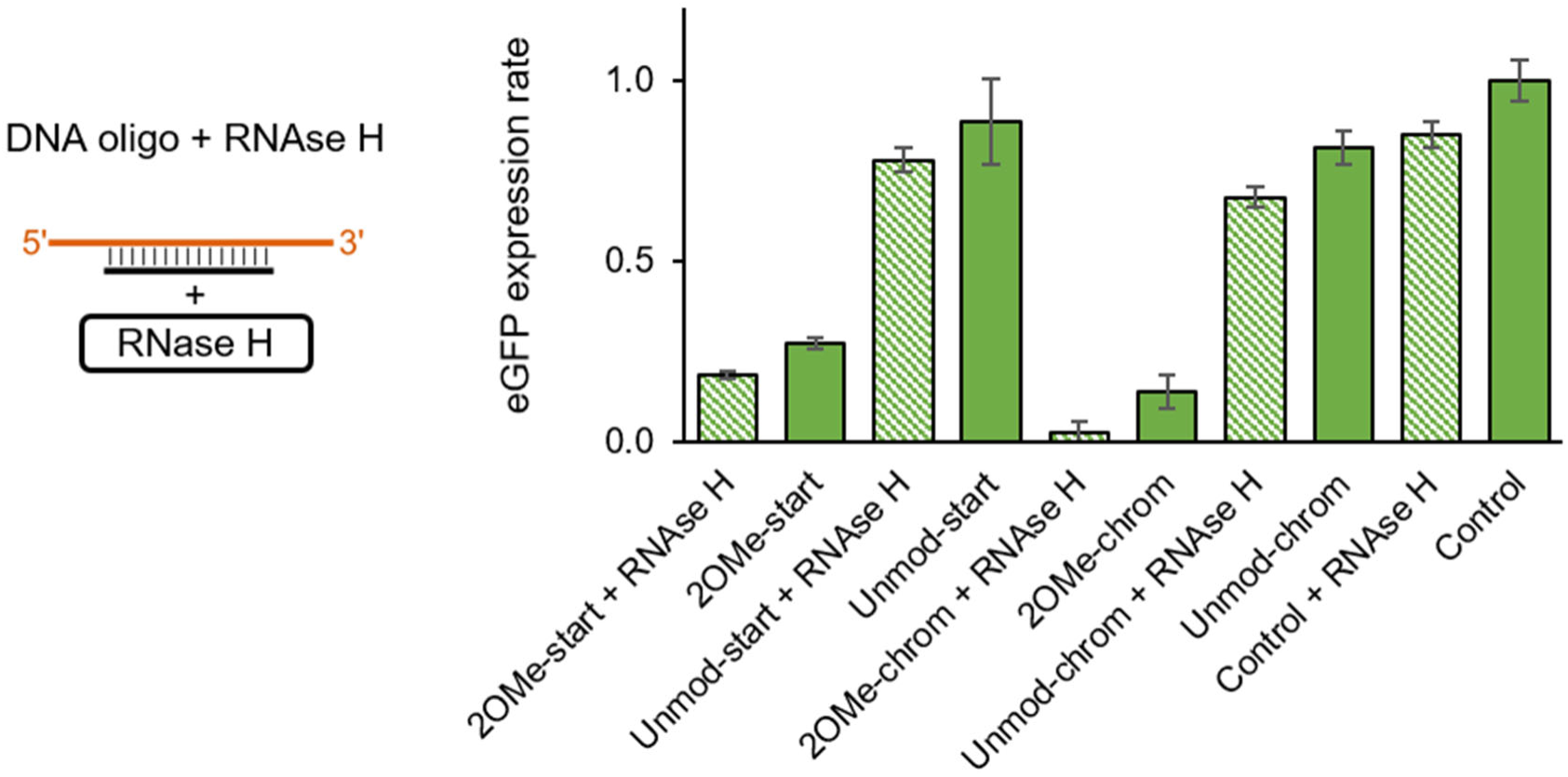
The effect of excess amount of RNase H supplementation in Akaby TXTL with silencing oligos. eGFP expression in Akaby TXTL with 2OMe or unmodified DNA oligos. 0.25 units/μl RNase H was added in the reactions indicated with “+ RNase H.” Control is the reaction with eGFP plasmid without oligo. Error bars indicate SEM, n=3.

**Supplementary Figure 10.**
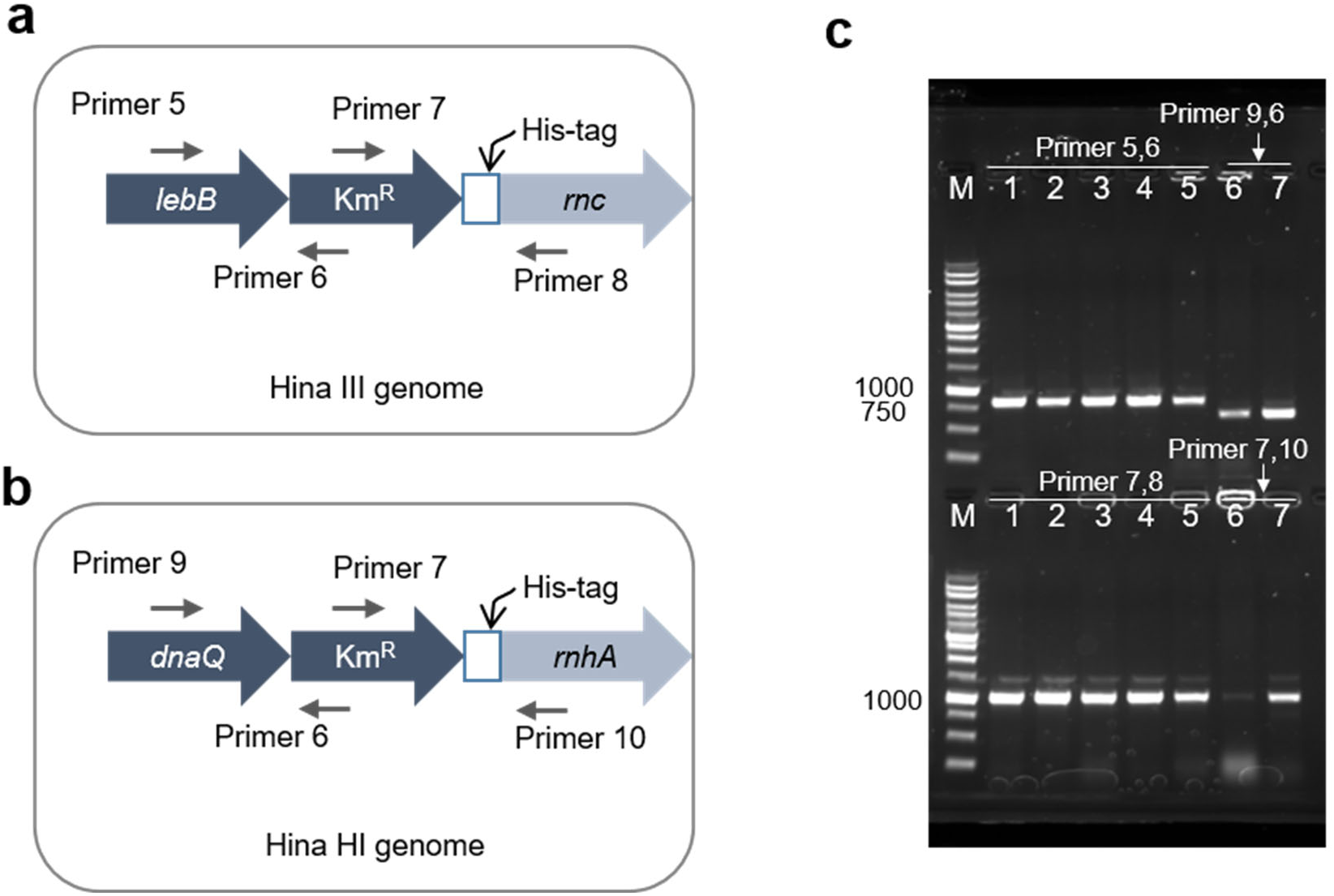
Colony PCR to confirm successful Hina III and Hina HI engineering. **a, b** The colony PCR primer designs for Hina III (**a**) and Hina HI (**b**) strains. Primer sequences are listed in **Supplementary Table 3**. Primer 6 and 7 are the insert specific primers, targeting kanamycin resistant gene (Km^R^.) Primer 5, 8, 9, and 10 are the locus-specific primers. RNase III and RNase HI are encoded by *rnc* and *rnhA* genes, respectively. **c** Agarose gel image of colony PCR. 1% agarose gel was run at 125 V for 20 minutes, stained with sYBR safe DNA Gel stain. Lane M: 1 kb DNA ladder (Goldbio, D010-500), Lane 1-5: Hina III colonies, Lane 6 and 7: Hina HI colonies, the expected product size: upper lane 1-5, 842 bp; upper lane 6 and 7, 666 bp; bottom lane 1-5, 1038 bp; bottom lane 6 and 7, 1042 bp.

**Supplementary Figure 11.**
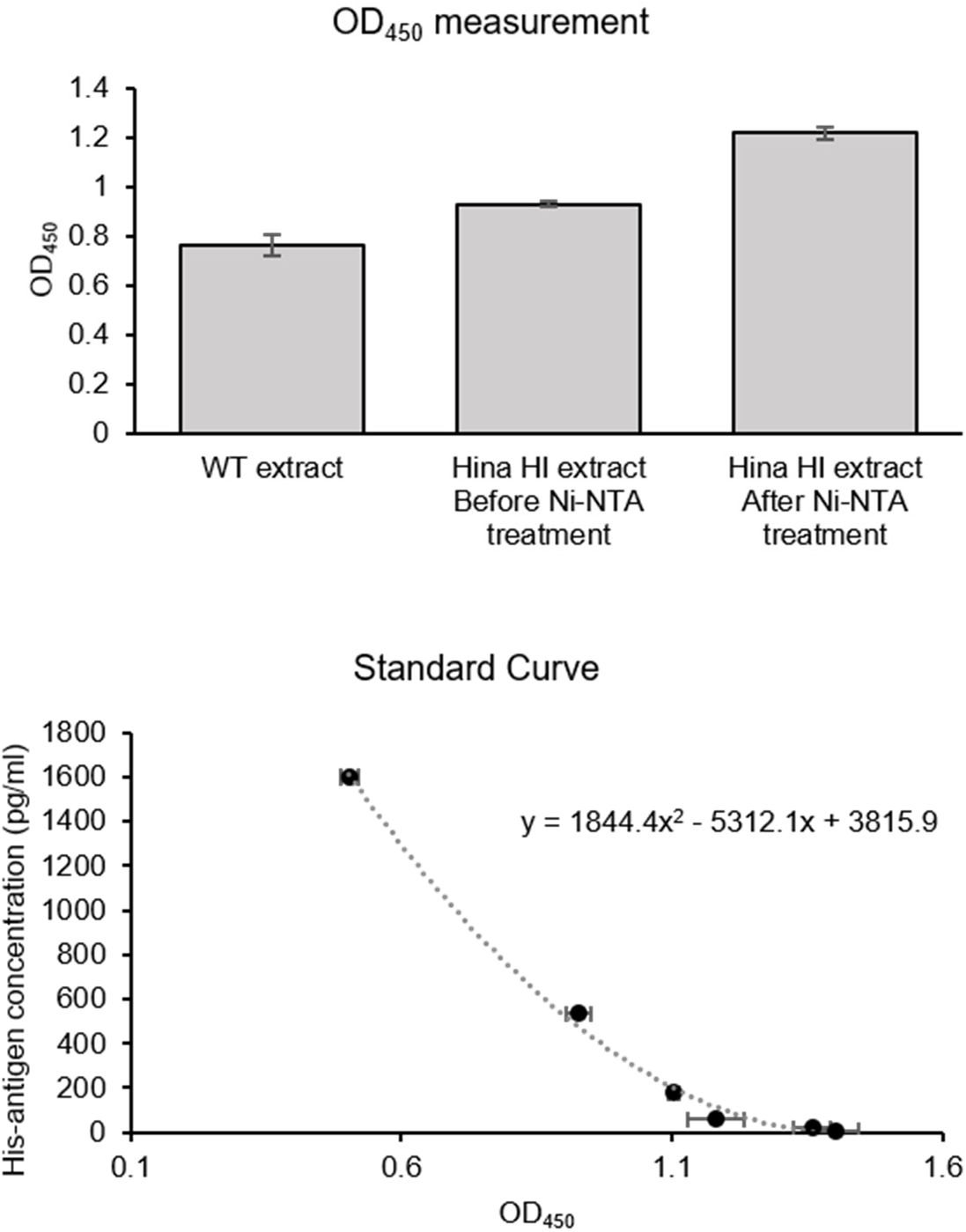
ELISA assay for detecting His-tagged proteins in TXTL. His Tag Elisa kit (LS-F55748) was purchased from LifeSpan Bioscience, inc. The cell-free preps were diluted to one-fifth with water, and the ELISA assay was performed following the manufacturer’s protocol. The standard curve was prepared using the standard sample prepared by the manufacturer. We confirmed the significant decrease of His-tagged protein in the Hina HI extract (After Ni-NTA treatment) compared to the Hina HI extract (Before Ni-NTA treatment. Error bars indicate SEM, n=3.

**Supplementary Figure 12.**
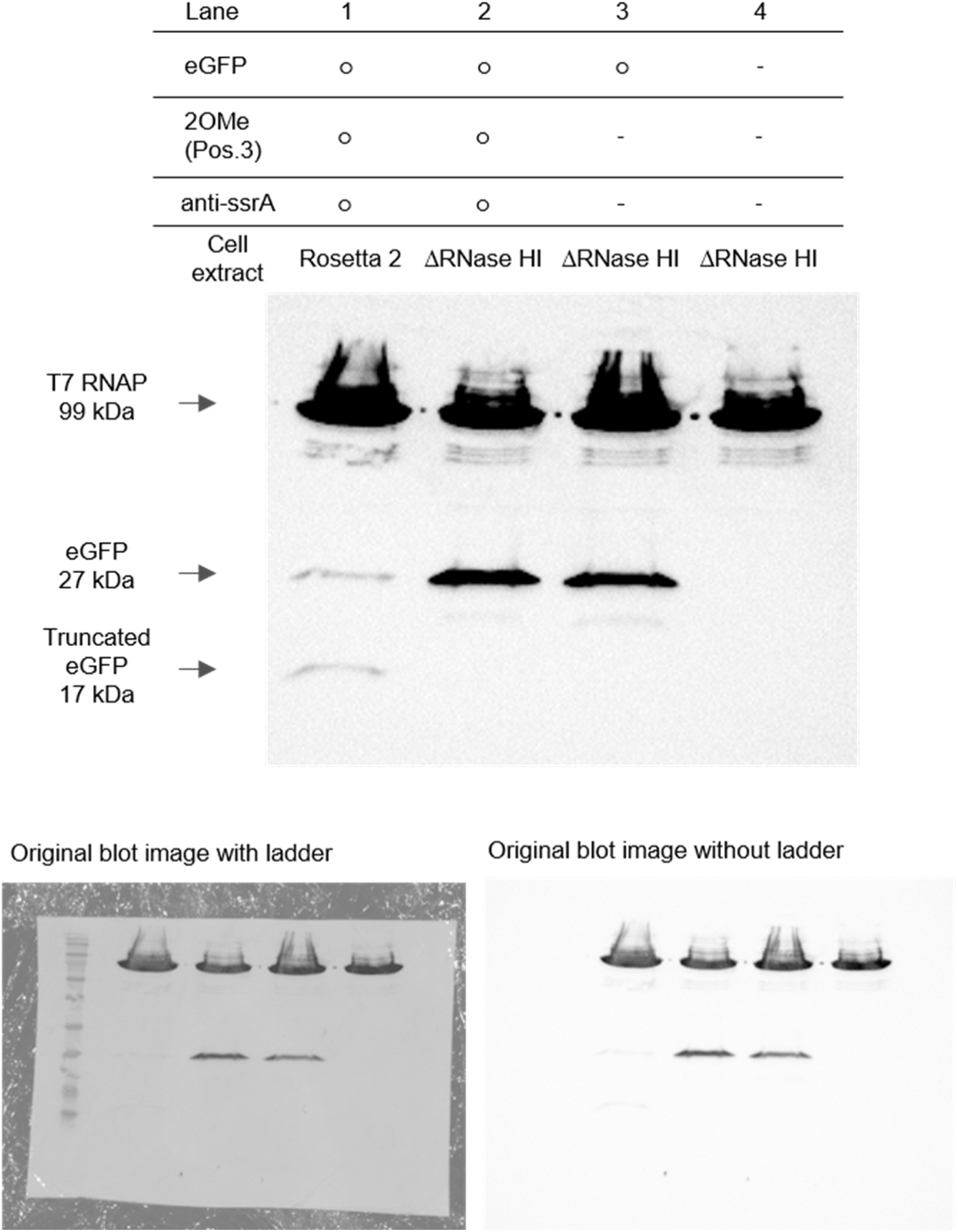
Western blotting of ΔRNase HI TXTL reactions. TXTL reactions were fractionated on a 12% polyacrylamide gel for 90 minutes. N-terminal His-tagged T7 RNA polymerase is 99 kDa. Full-length N-terminal His-tagged eGFP is 27 kDa. The truncated eGFP by 2OMe-oligo interruption is 17 kDa. anti-ssrA oligo is added to prevent tmRNA-associated truncated protein degradation. The table above the gel indicates which components mixed in the TXTL reaction: “○” means the component was contained in the reaction and “ – ” means the component was not added. While normal TXTL (Rosetta 2) shows the truncated eGFP, ΔRNase HI TXTL did not produce it even with anti-ssrA supplementation.

**Supplementary Figure 13.**
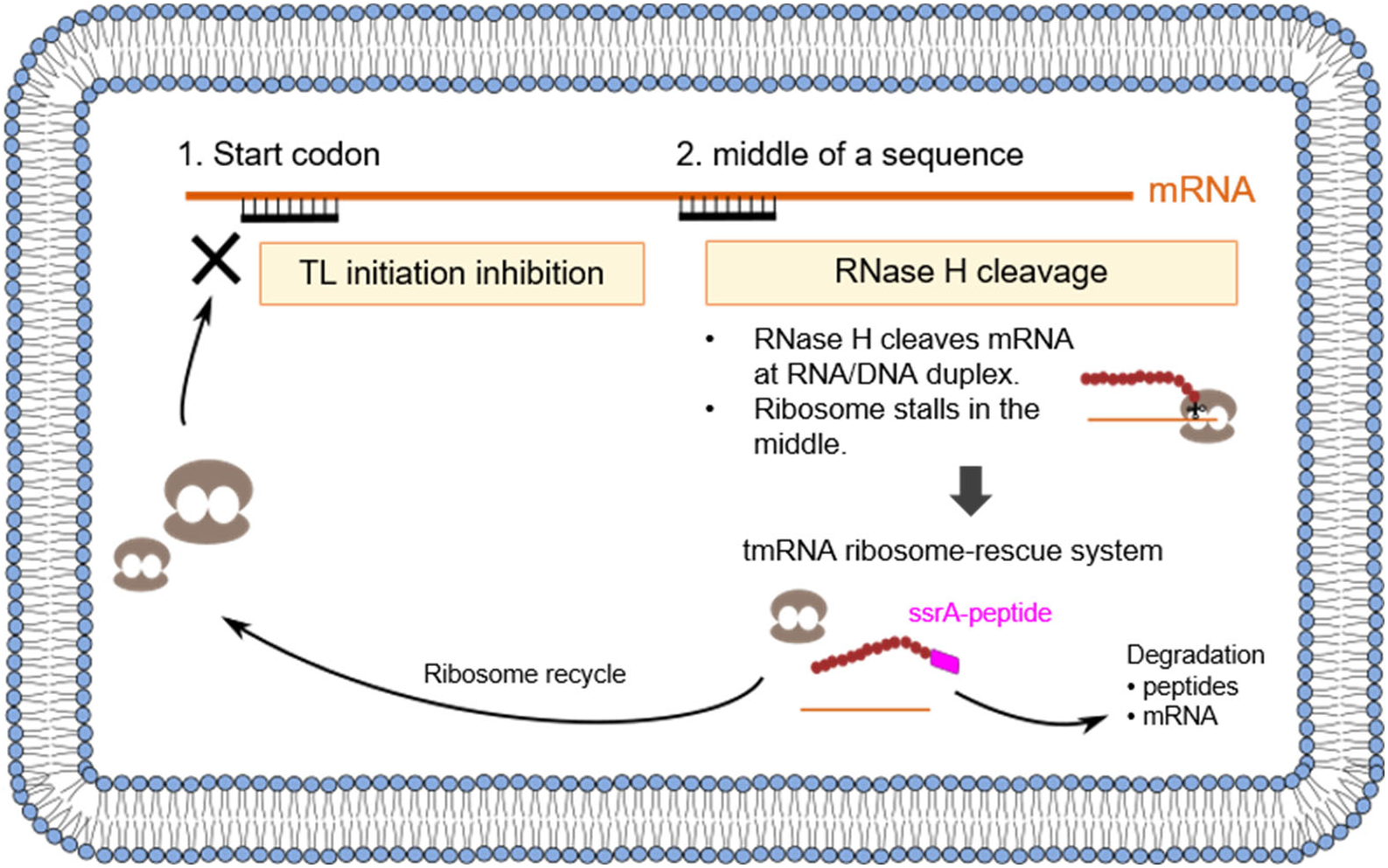
Gene silencing mechanism of 2OMe-oligo. 2OMe-oligo inhibits gene expression in two different mechanisms. When 2OMe-oligo binds near the start codon, the gene expression is inhibited by preventing ribosomes from starting translation. When 2OMe-oligo binds somewhere in the middle of the sequence, RNase H cleaves the mRNA at the DNA/RNA duplex. Ribosomes translate the mRNA up to the cleaved mRNA position, then get stalled. tmRNA releases the stalled ribosomes and adds a ssrA-tag on the partially translated protein. The mRNA and the truncated protein are degraded, and ribosomes are recycled.

**Supplementary Figure 14.**
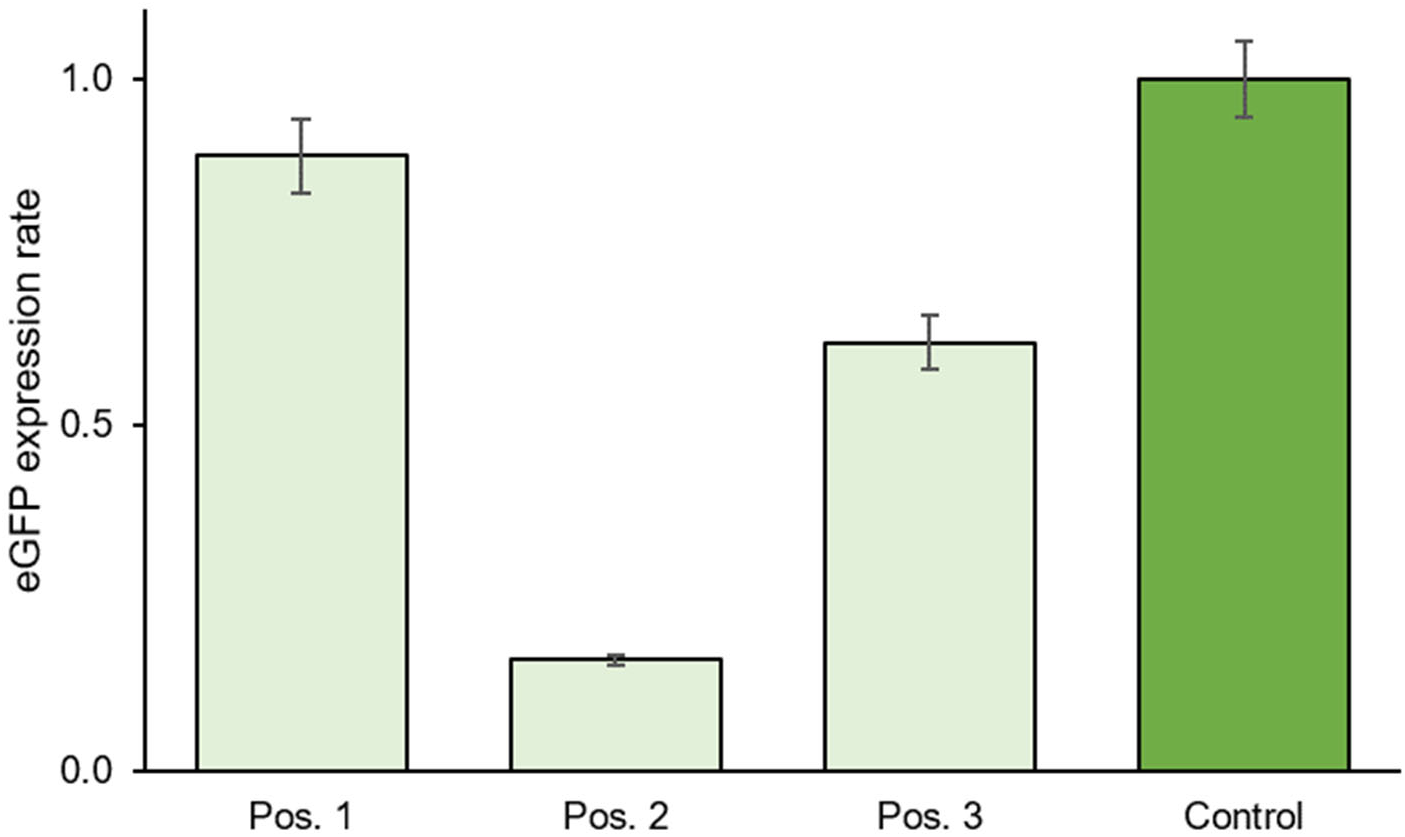
2OMe-oligo inhibition in eGFP expression in ΔRNase III TXTL. eGFP expression was measured in ΔRNase III TXTL with 2OMe-oligo, Pos.1, Pos.2, and Pos.3. Control is the reaction with eGFP plasmid without 2OMe-oligo. Error bars indicate SEM, n=3.

**Supplementary Figure 15.**
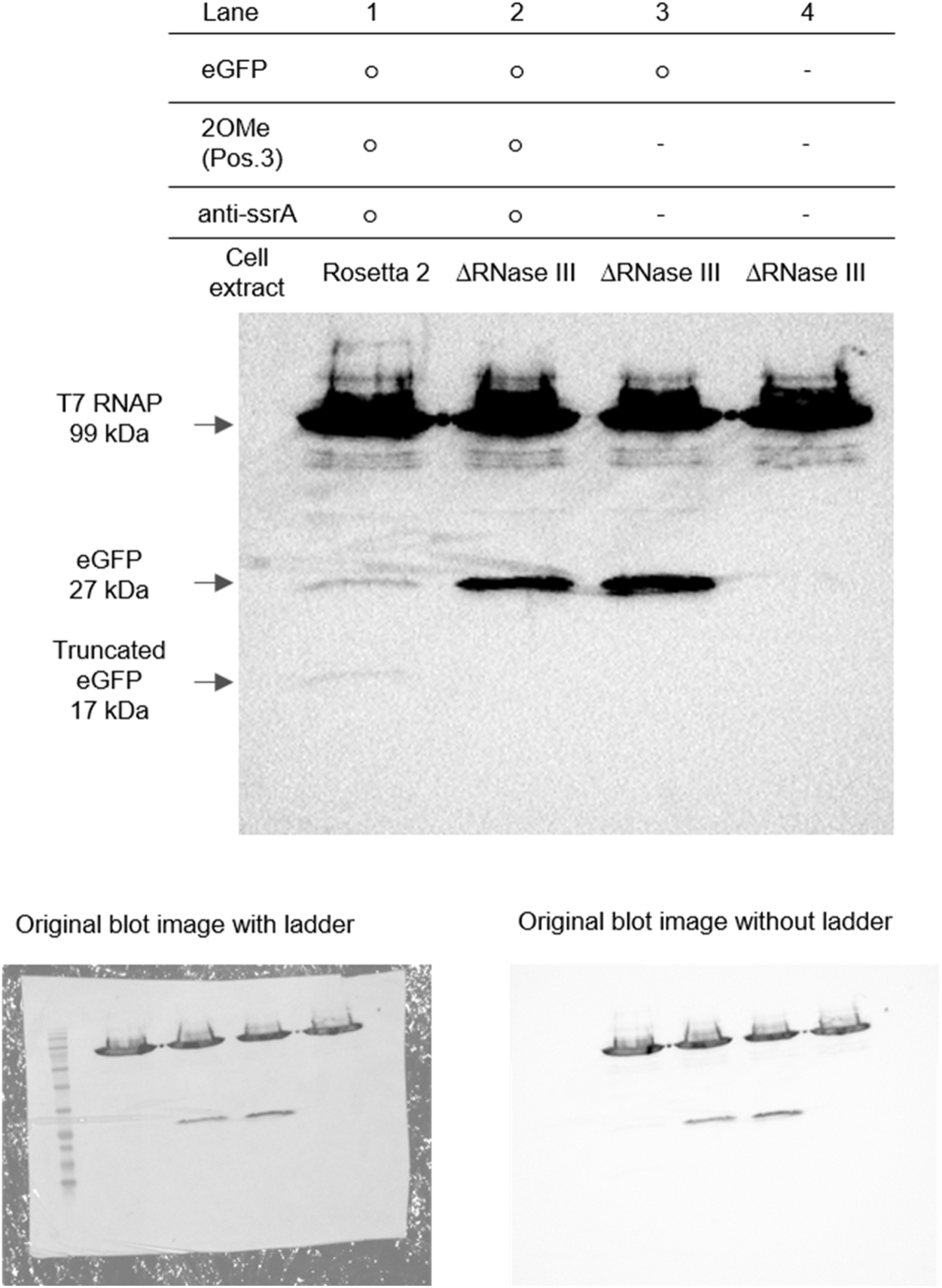
Western blotting image of eGFP expression in ΔRNase III TXTL with anti-ssrA oligo. TXTL reactions were fractionated on a 12% polyacrylamide gel for 90 minutes. N-terminal His-tagged T7 RNA polymerase is 99 kDa. Full-length N-terminal His-tagged eGFP is 27 kDa. The truncated eGFP by 2OMe-oligo interruption is 17 kDa. anti-ssrA oligo is added to prevent tmRNA-associated truncated protein degradation. The table above the gel indicates which components mixed in the TXTL reaction: “○” means the component was contained in the reaction and “ – ” means the component was not added. While normal TXTL (Rosetta 2) shows the truncated eGFP, ΔRNase III TXTL did not produce it even with anti-ssrA supplementation.

**Supplementary Figure 16.**
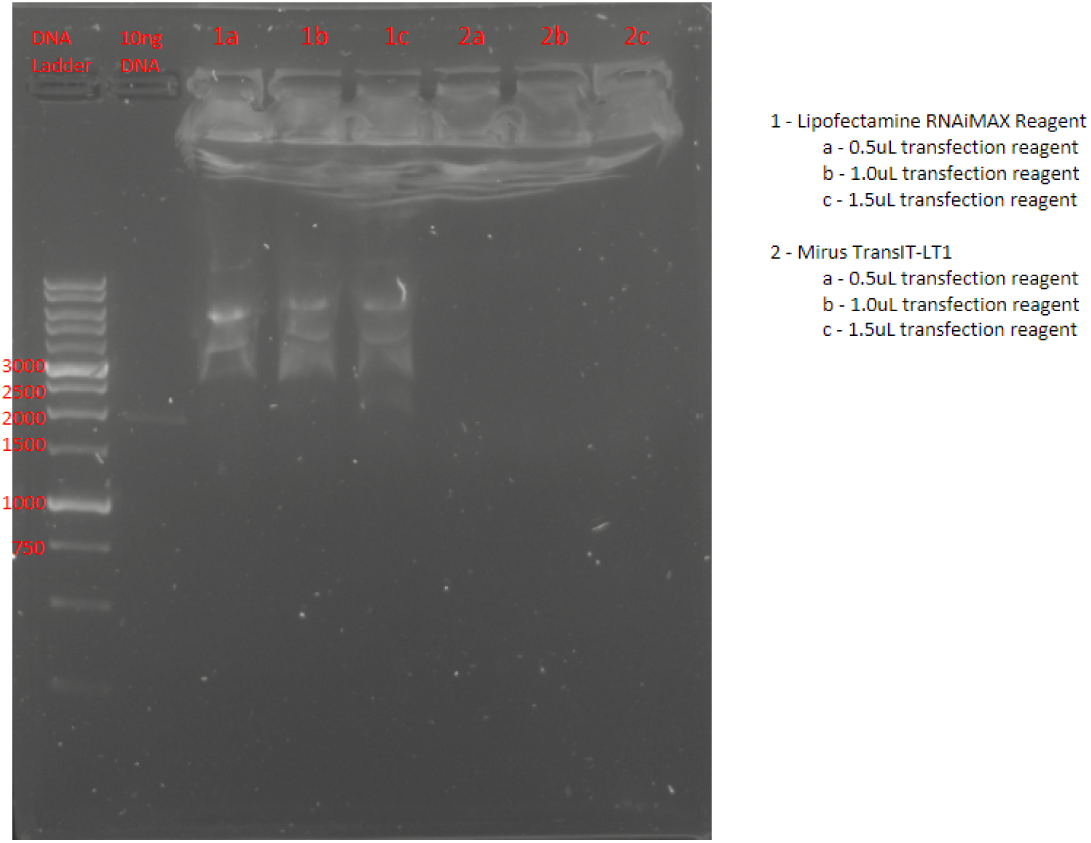
Analysis of restriction enzyme digest of transfection reagent test in synthetic cell liposomes. Samples on 1% agarose gel with SybrSafe.

**Supplementary Figure 17.**
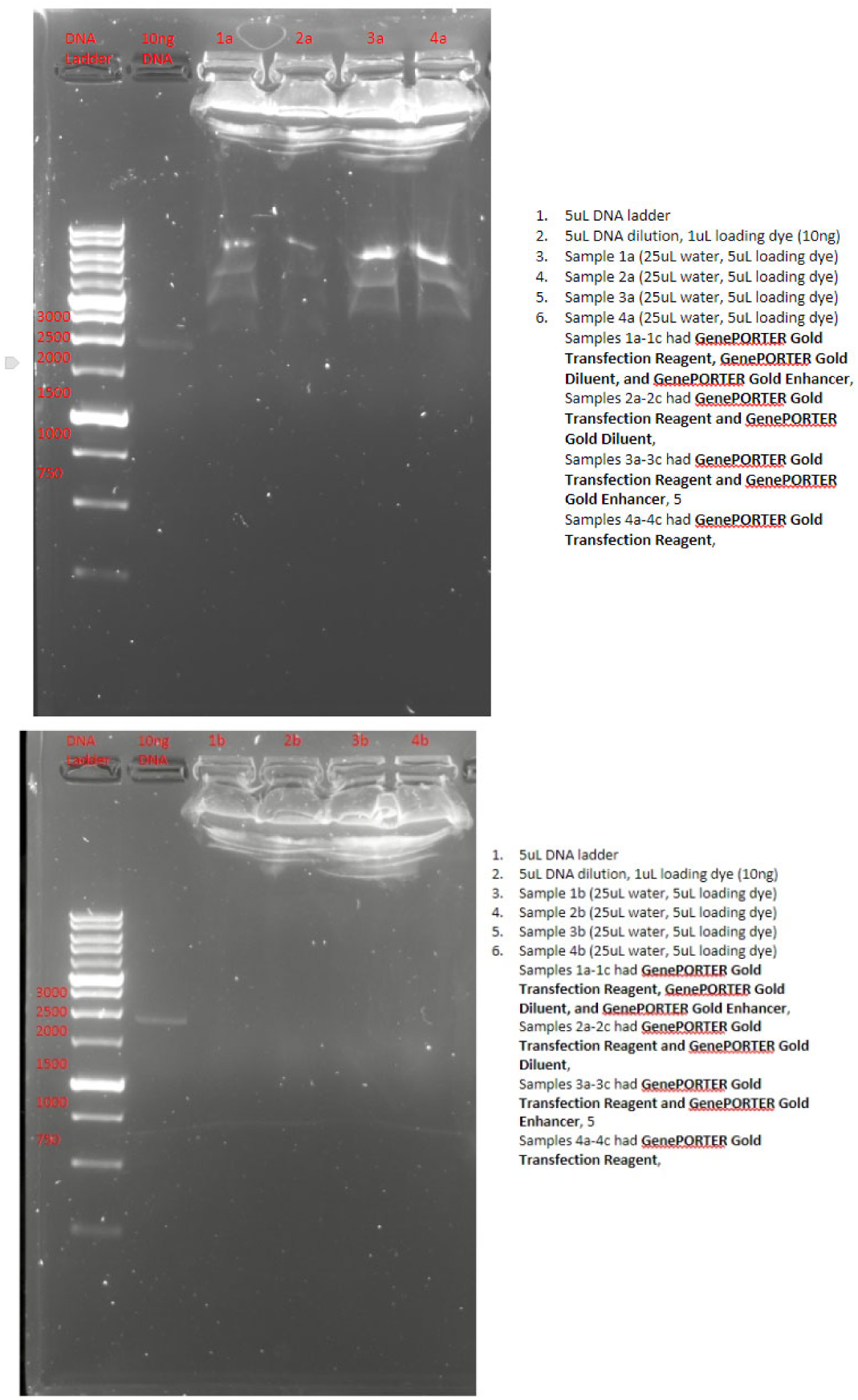
Analysis of restriction enzyme digest of transfection reagent test in synthetic cell liposomes. Samples on 1% agarose gel with SybrSafe.

**Supplementary Figure 18.**
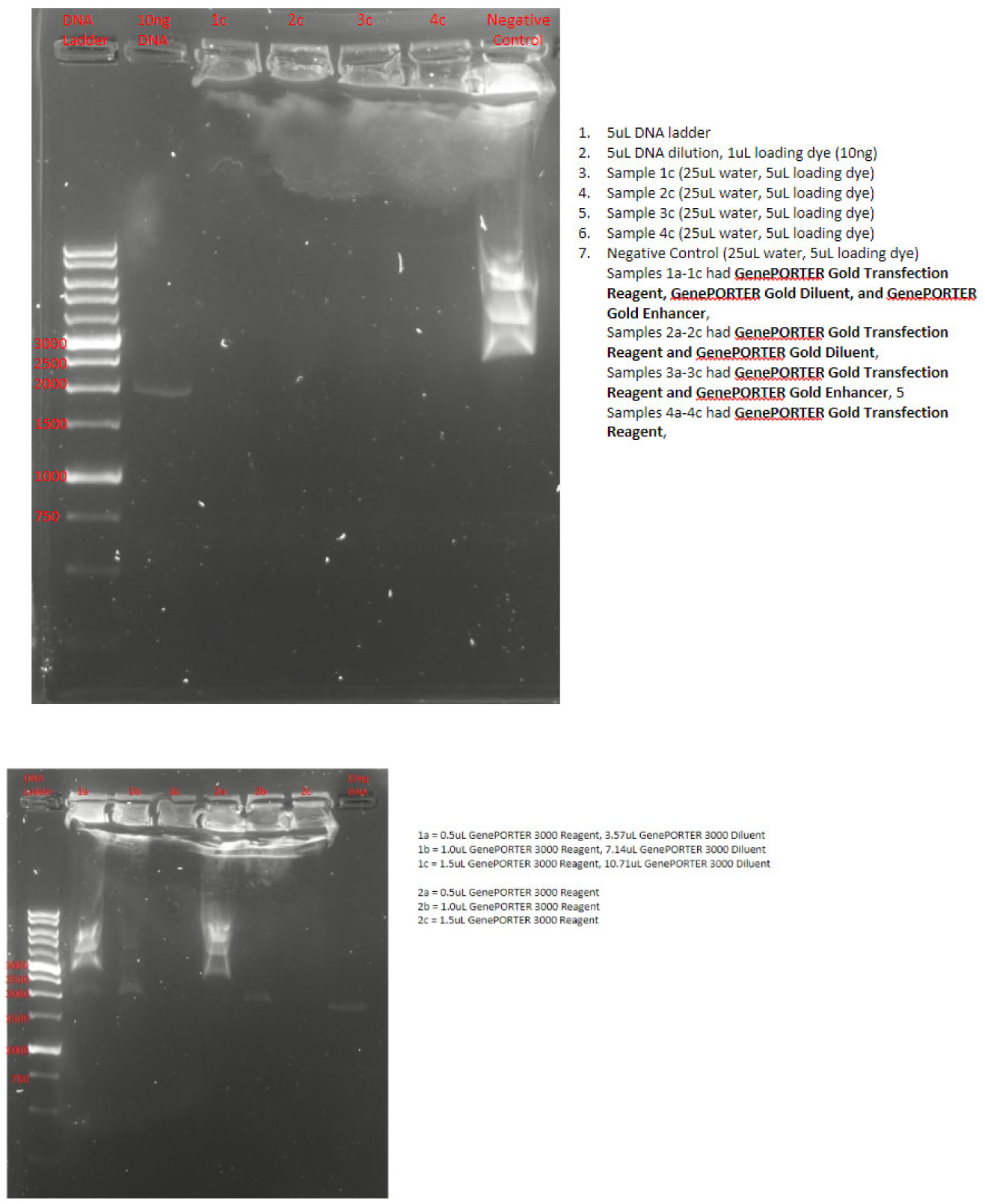
Analysis of restriction enzyme digest of transfection reagent test in synthetic cell liposomes. Samples on 1% agarose gel with SybrSafe.

**Supplementary Figure 19.**
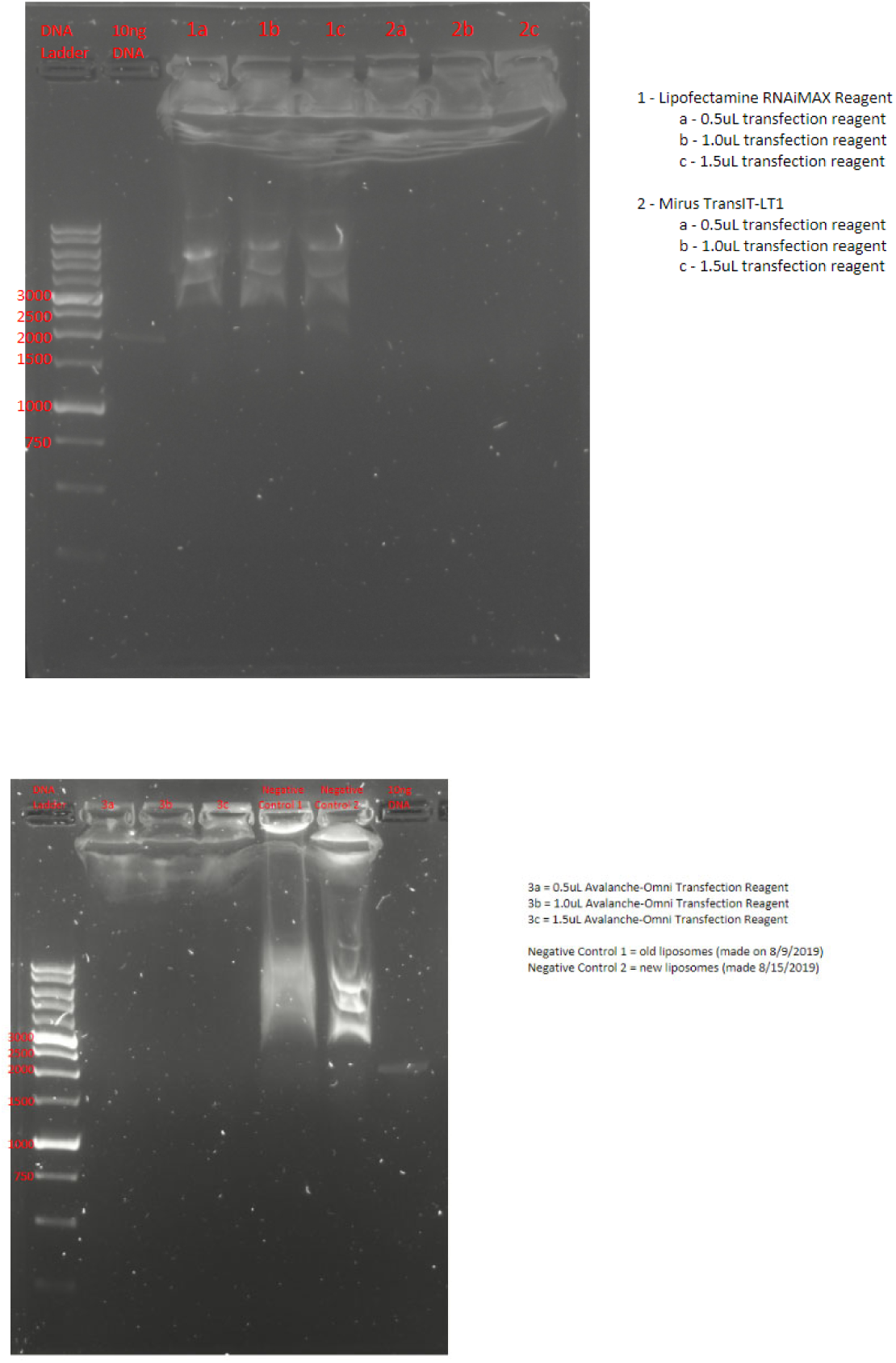
Analysis of restriction enzyme digest of transfection reagent test in synthetic cell liposomes. Samples on 1% agarose gel with SybrSafe.

**Supplementary Figure 20.**
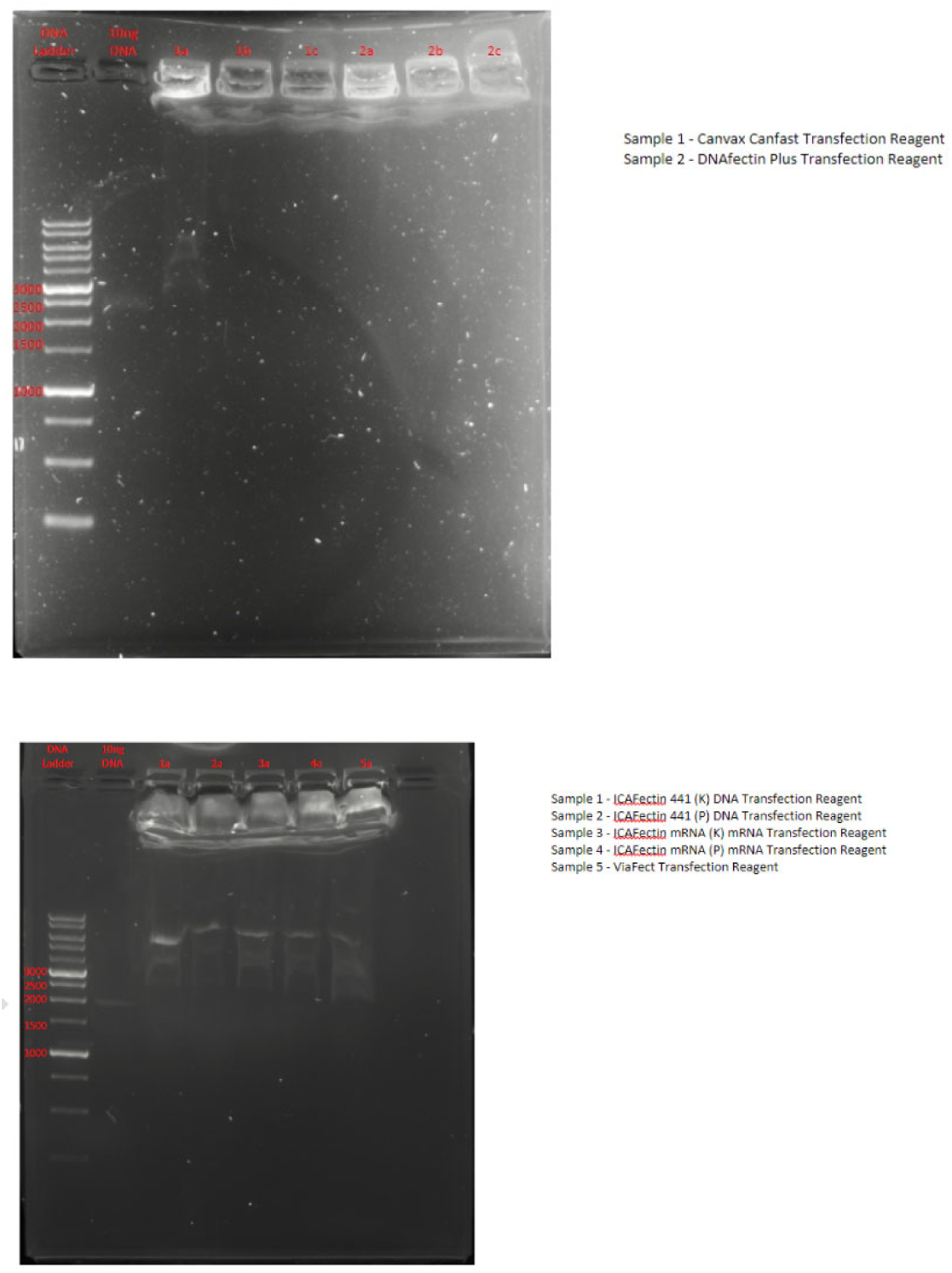
Analysis of restriction enzyme digest of transfection reagent test in synthetic cell liposomes. Samples on 1% agarose gel with SybrSafe.

**Supplementary Figure 21.**
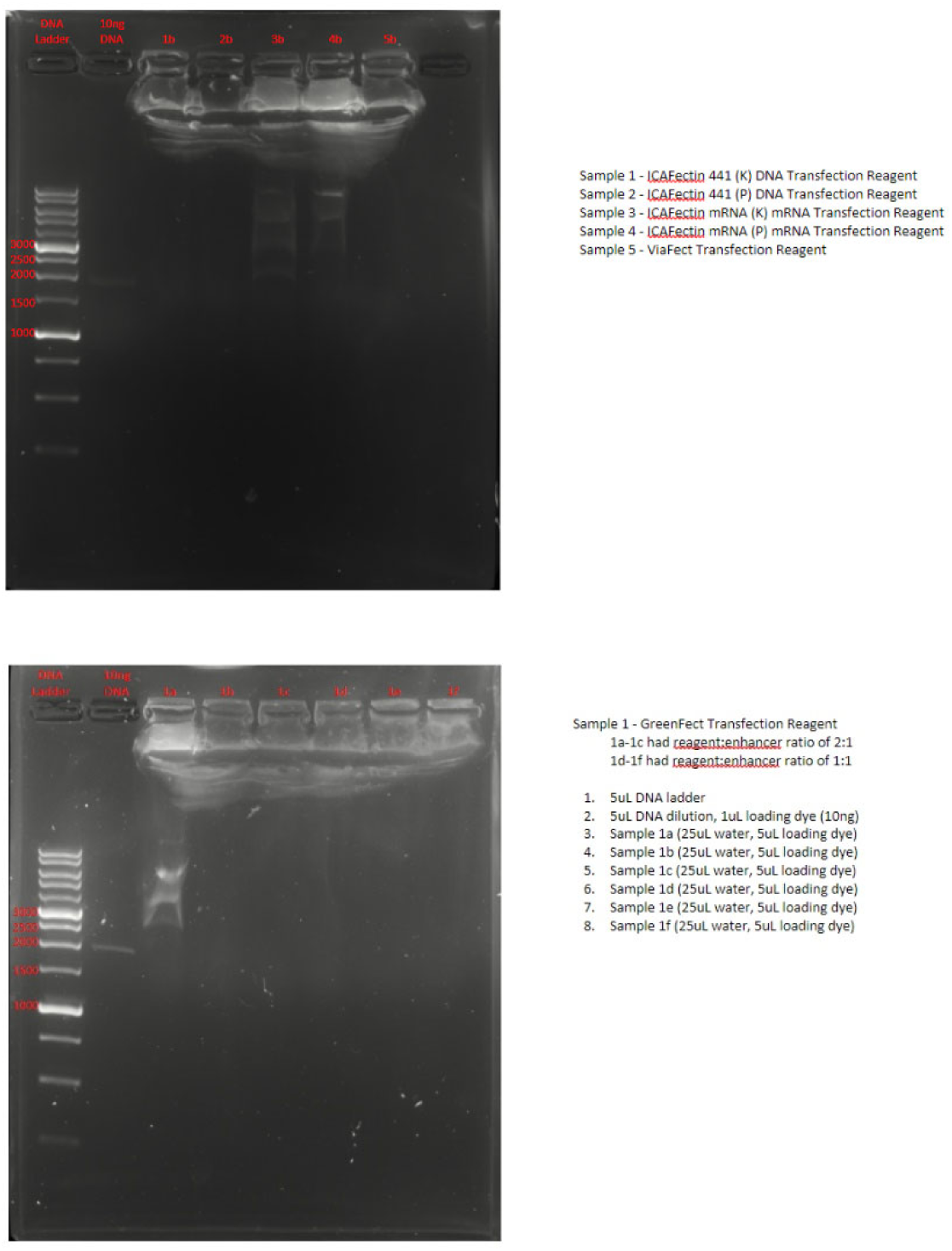
Analysis of restriction enzyme digest of transfection reagent test in synthetic cell liposomes. Samples on 1% agarose gel with SybrSafe.

**Supplementary Figure 22.**
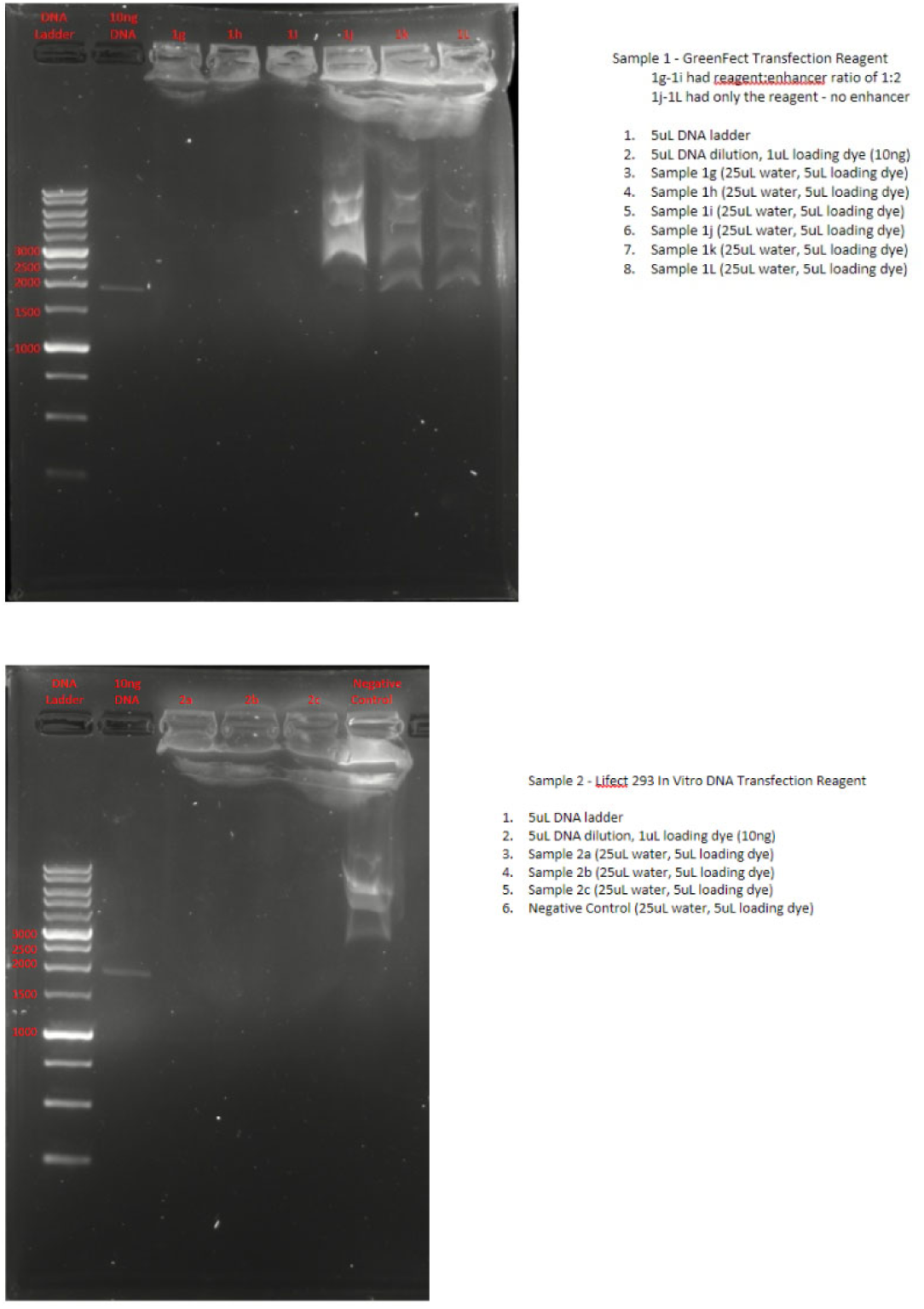
Analysis of restriction enzyme digest of transfection reagent test in synthetic cell liposomes. Samples on 1% agarose gel with SybrSafe.

**Supplementary Figure 23.**
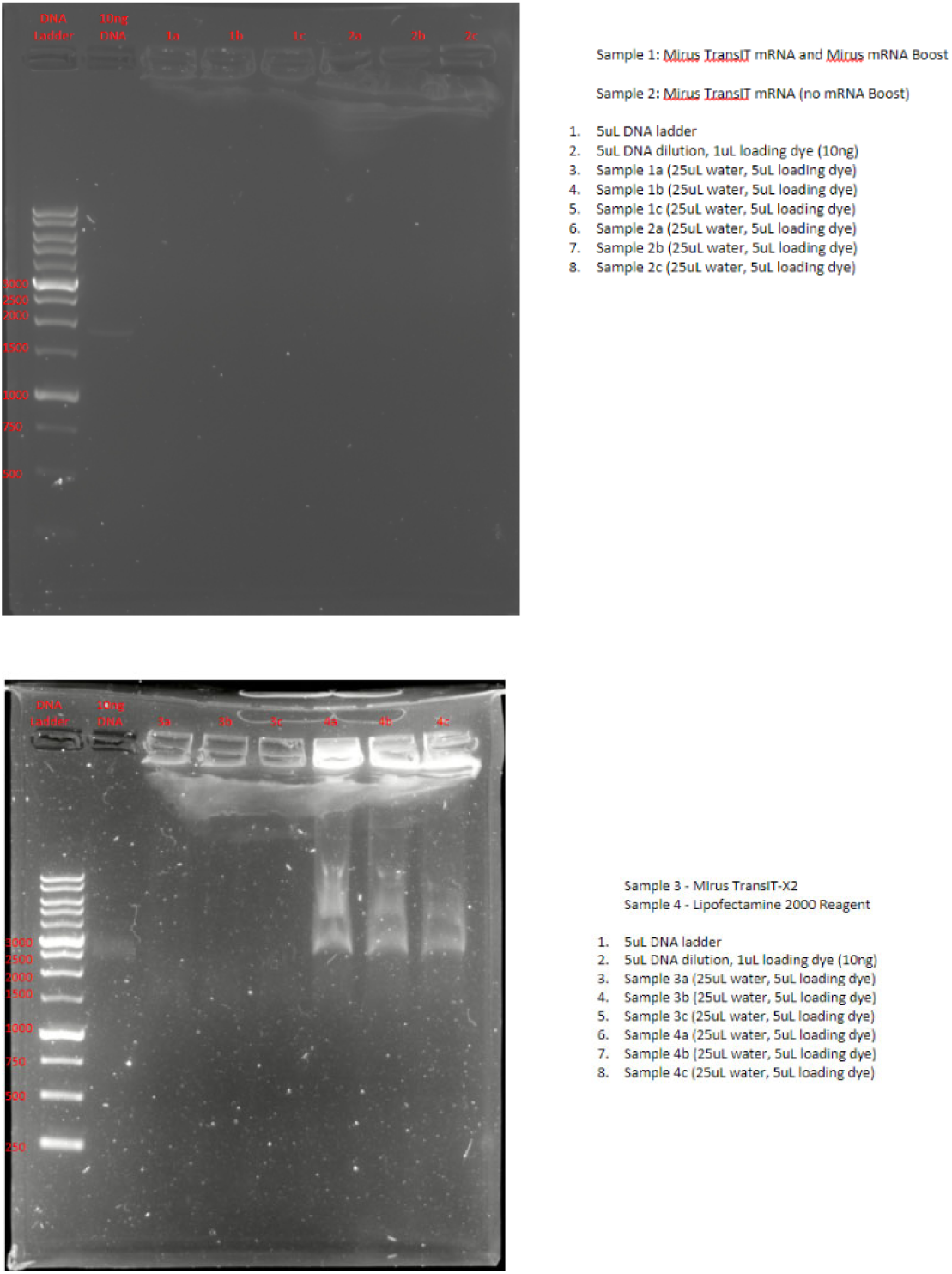
Analysis of restriction enzyme digest of transfection reagent test in synthetic cell liposomes. Samples on 1% agarose gel with SybrSafe.

**Supplementary Figure 24.**
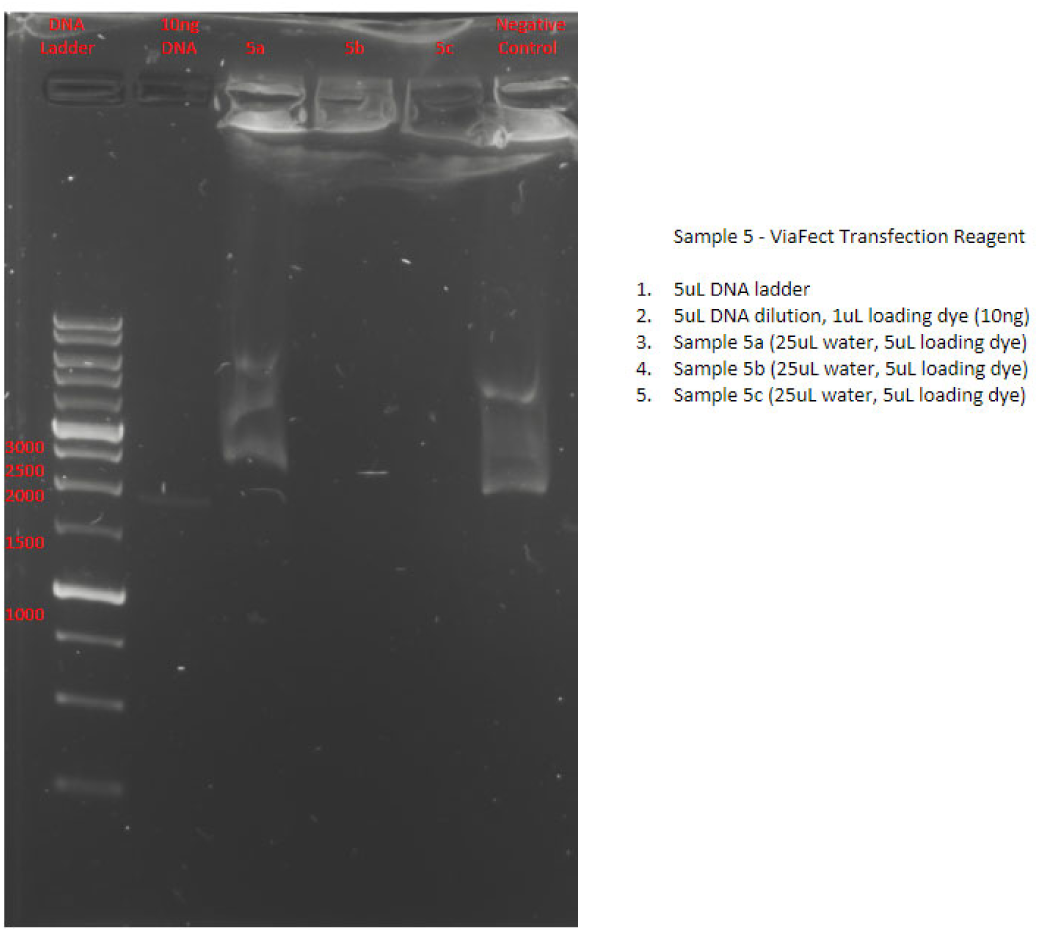
Analysis of restriction enzyme digest of transfection reagent test in synthetic cell liposomes. Samples on 1% agarose gel with SybrSafe.

**Supplementary Figure 25.**
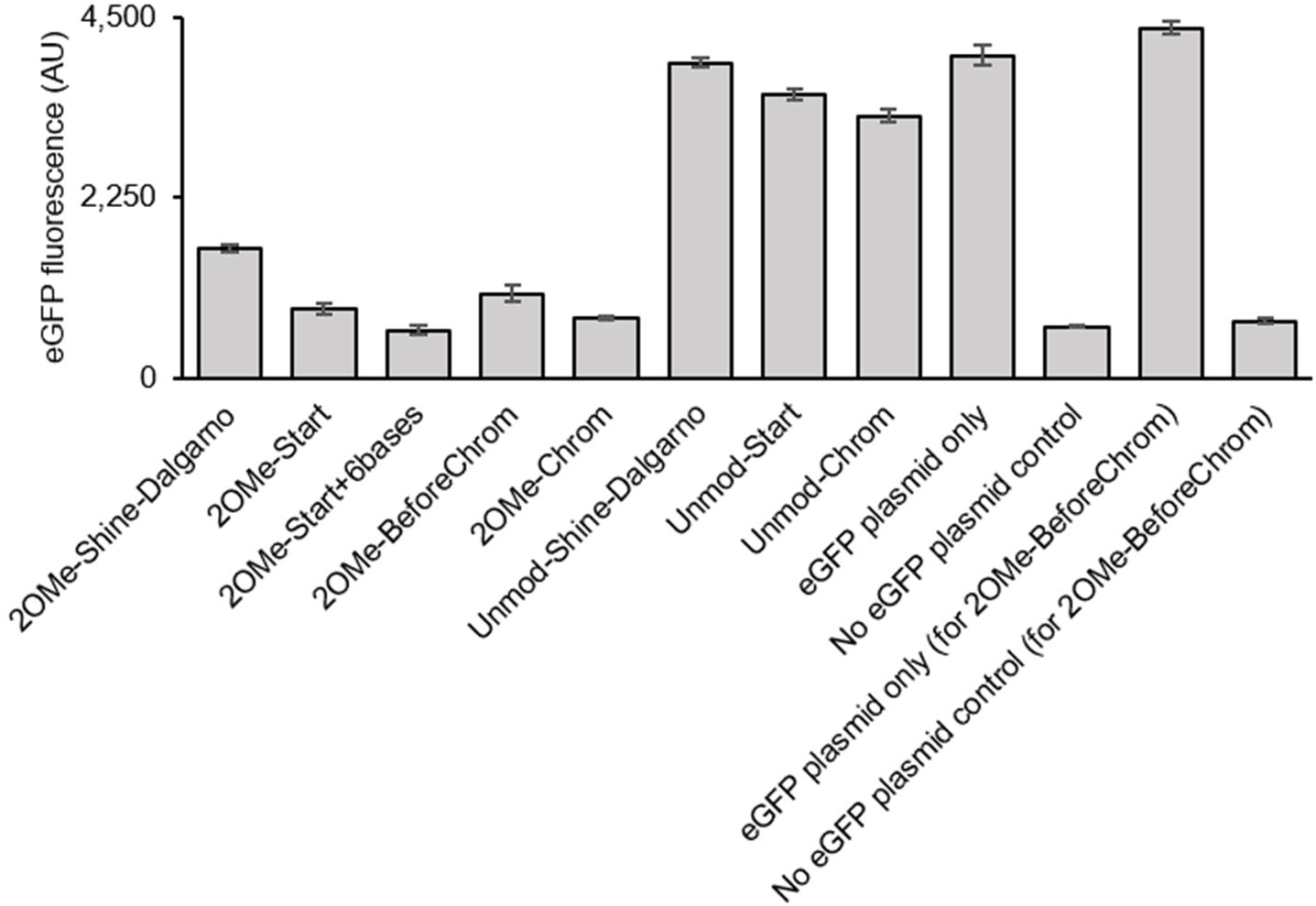
eGFP fluorescence of TXTL reactions shown in Fig. 1d.

**Supplementary Figure 26.**
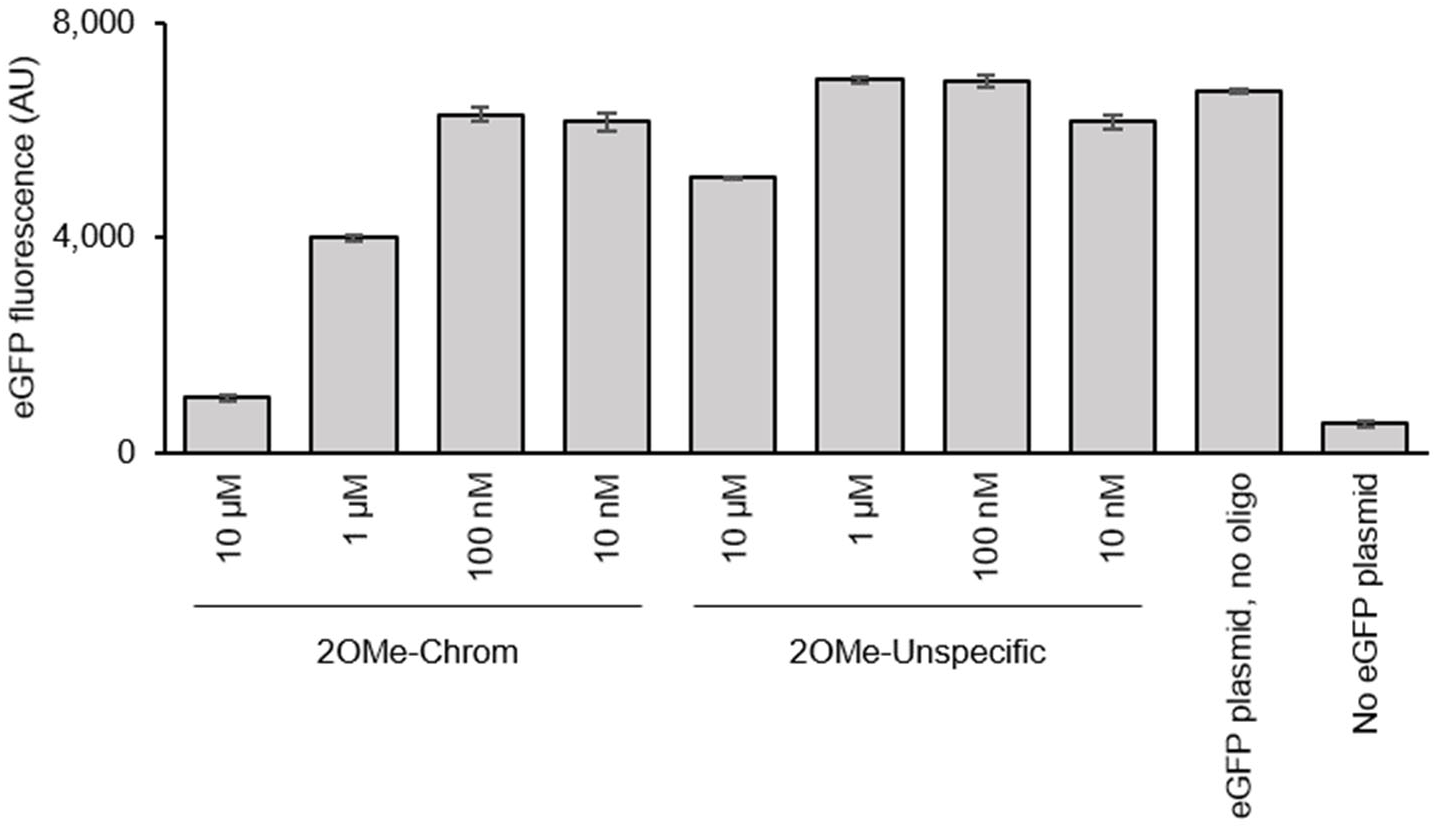
eGFP fluorescence of TXTL reactions shown in Fig. 1e.

**Supplementary Figure 27.**
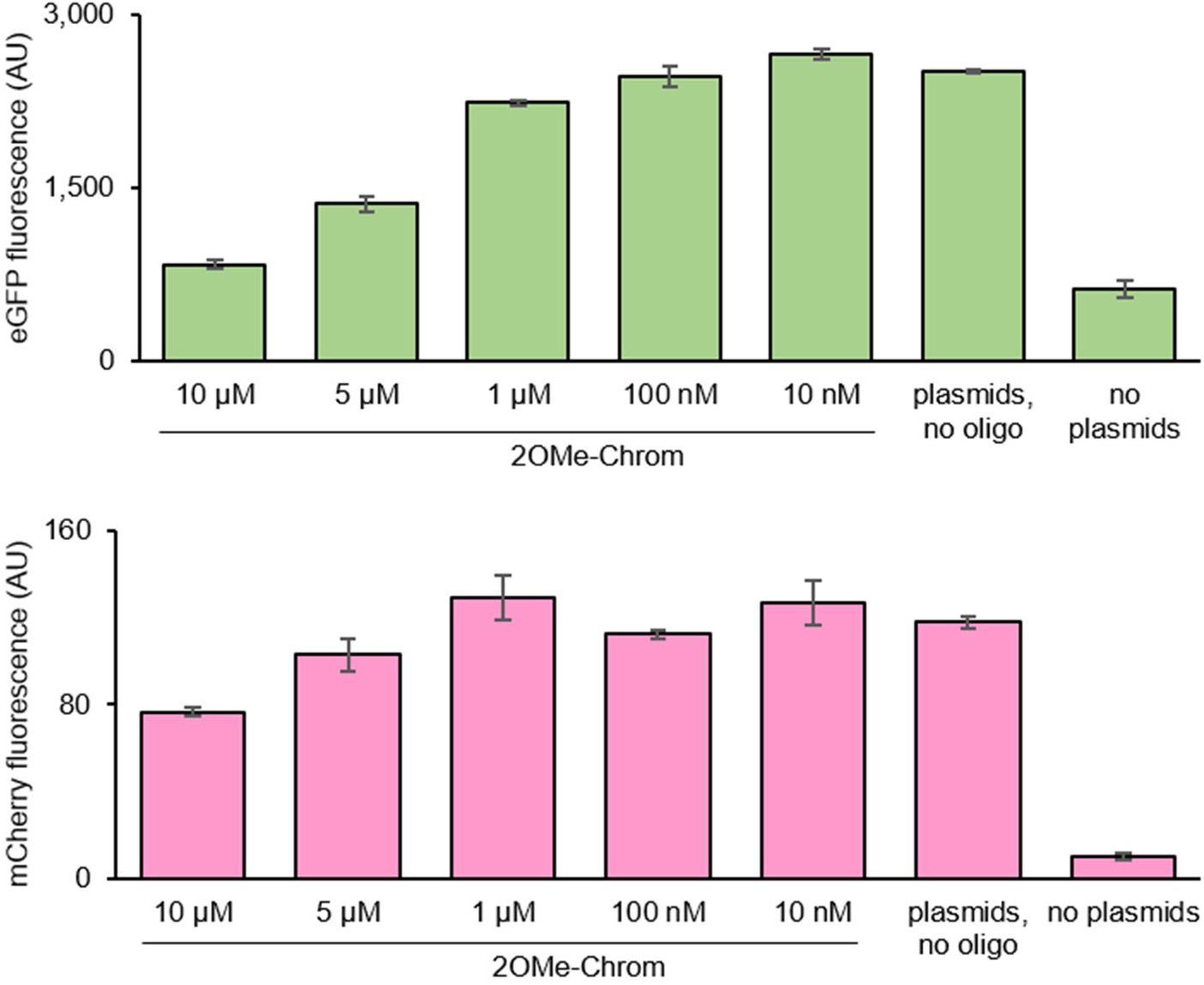
eGFP and mCherry fluorescence of TXTL reactions shown in Fig. 1f.

**Supplementary Figure 28.**
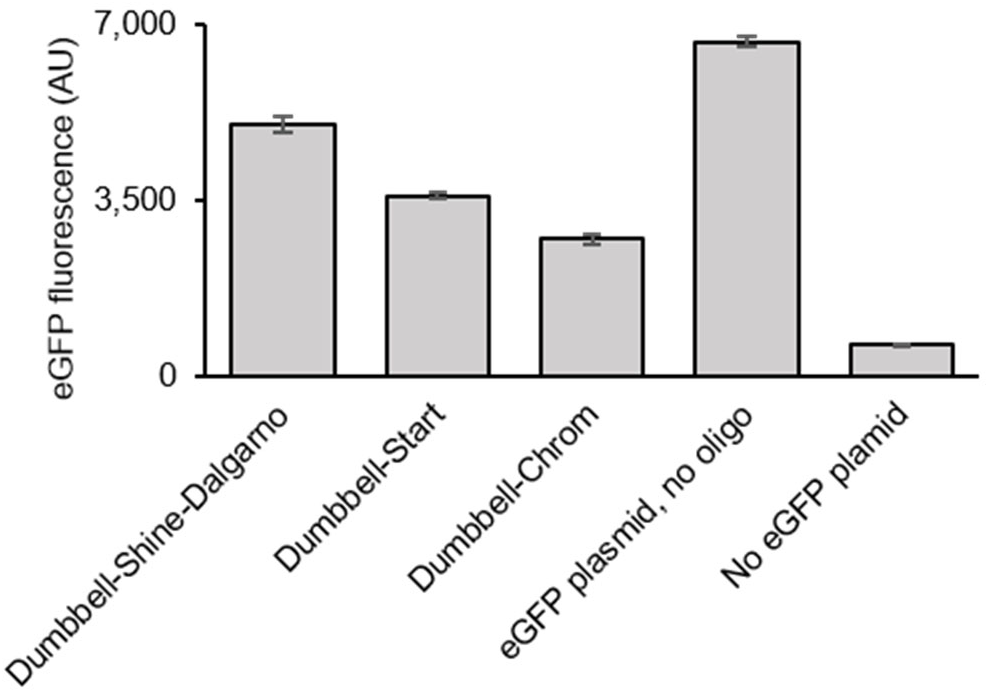
eGFP fluorescence of TXTL reactions shown in Fig. 1h.

**Supplementary Figure 29.**
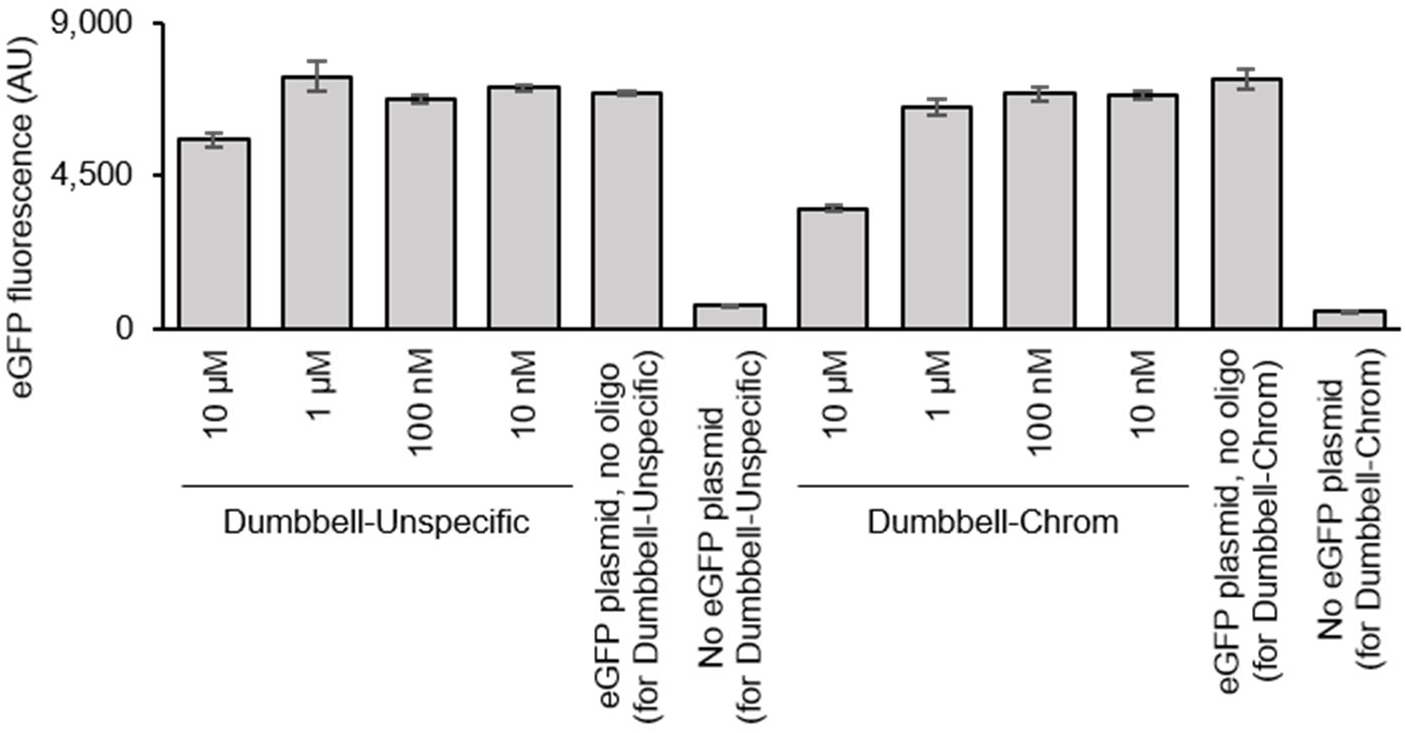
eGFP fluorescence of TXTL reactions shown in Fig. 1i.

**Supplementary Figure 30.**
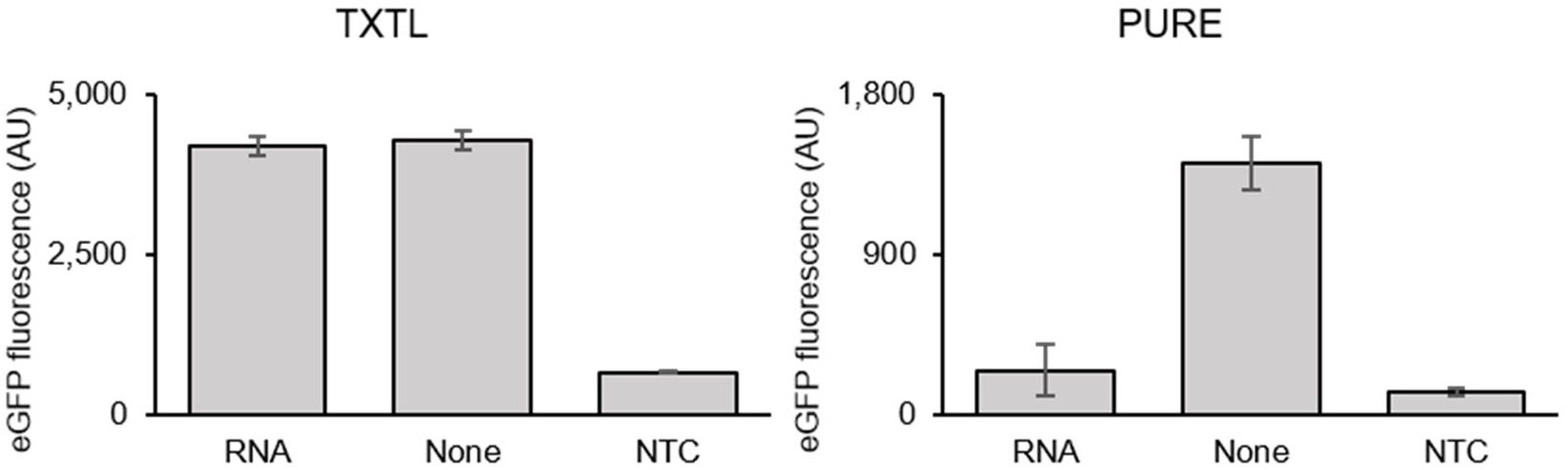
eGFP fluorescence of TXTL reactions shown in Fig. 1k.

**Supplementary Figure 31.**
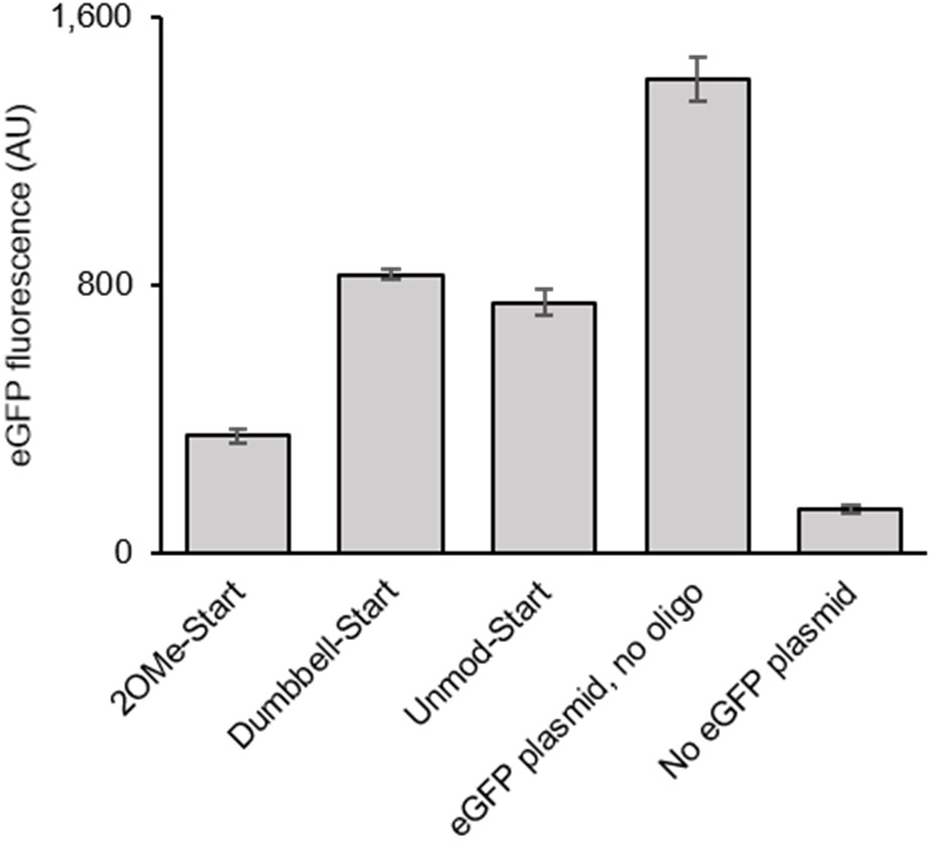
eGFP fluorescence of TXTL reactions shown in Fig. 1l.

**Supplementary Figure 32.**
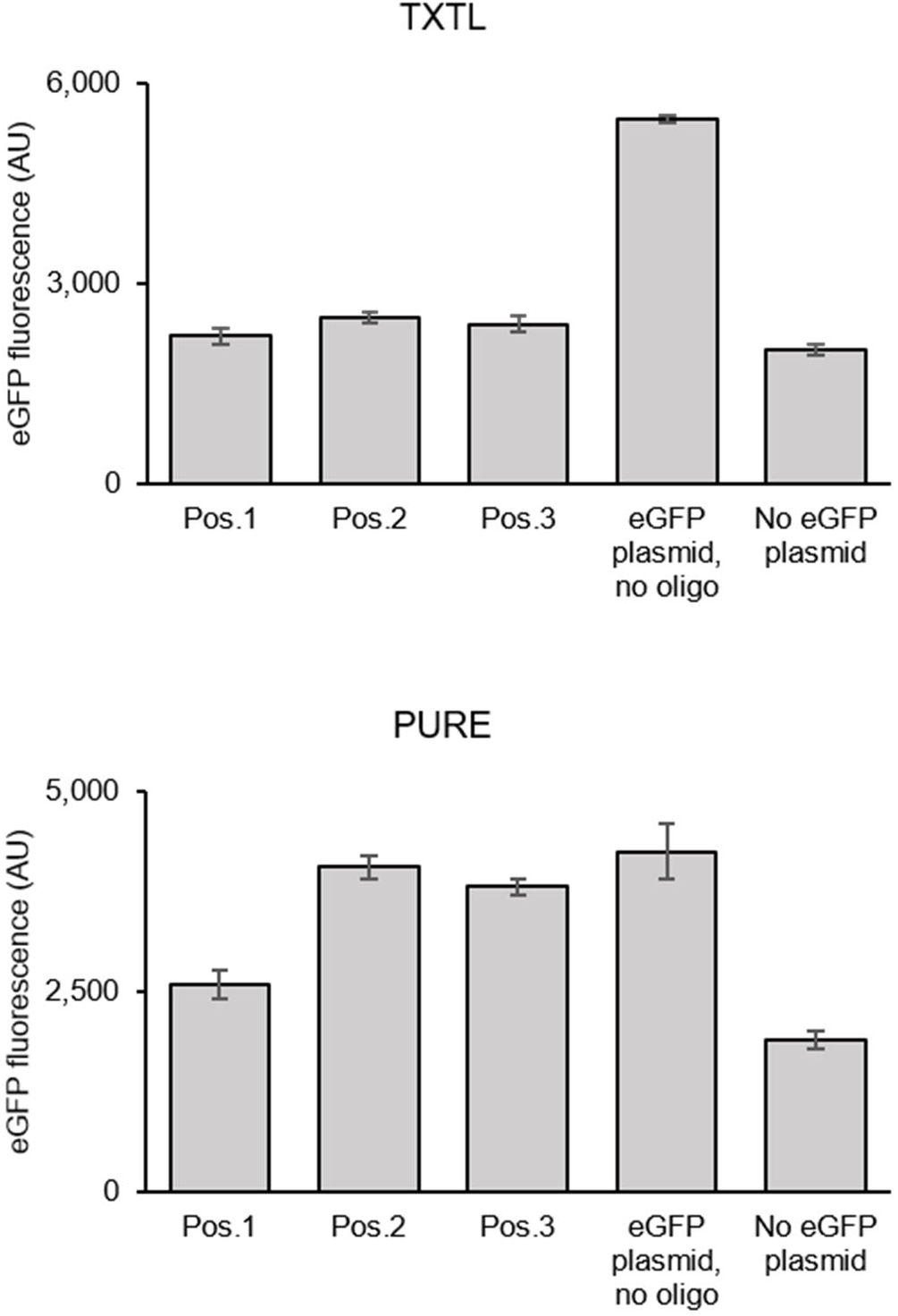
eGFP fluorescence of TXTL reactions shown in Fig. 2b and d.

**Supplementary Figure 33.**
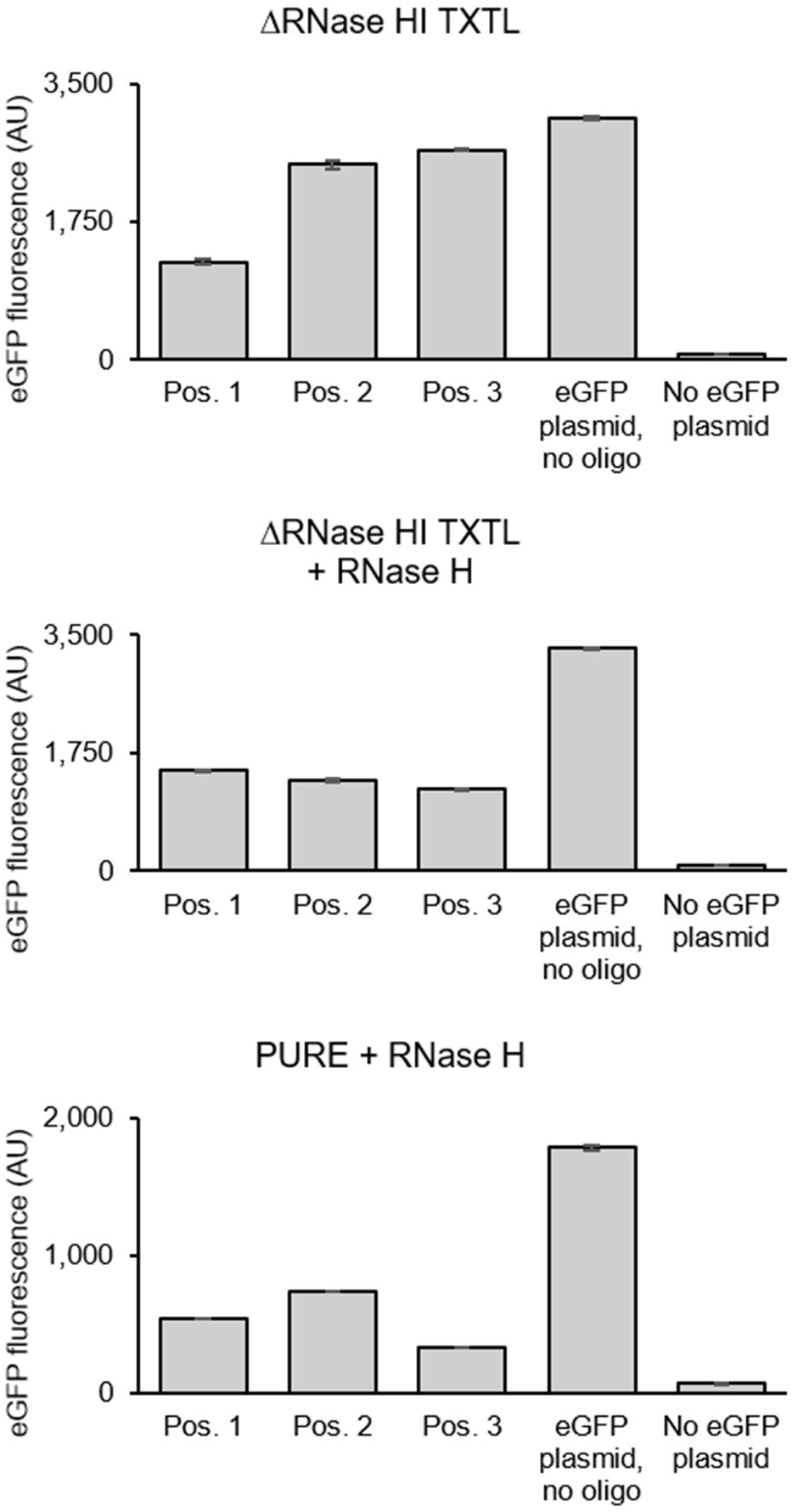
eGFP fluorescence of TXTL reactions shown in Fig. 2f, g, and h.

**Supplementary Figure 34.**
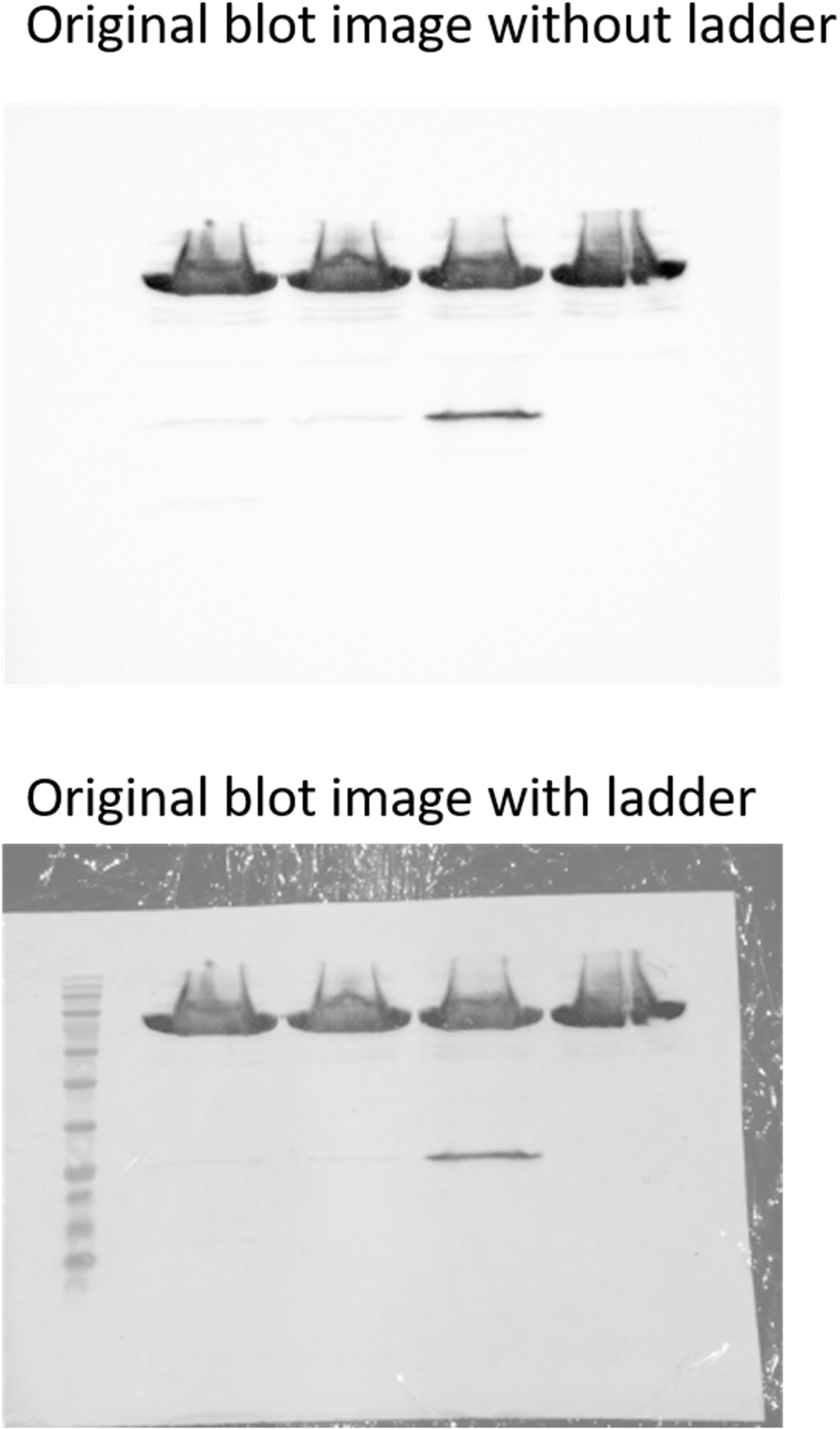
Original blot image shown in Fig. 2i.

**Supplementary Figure 35.**
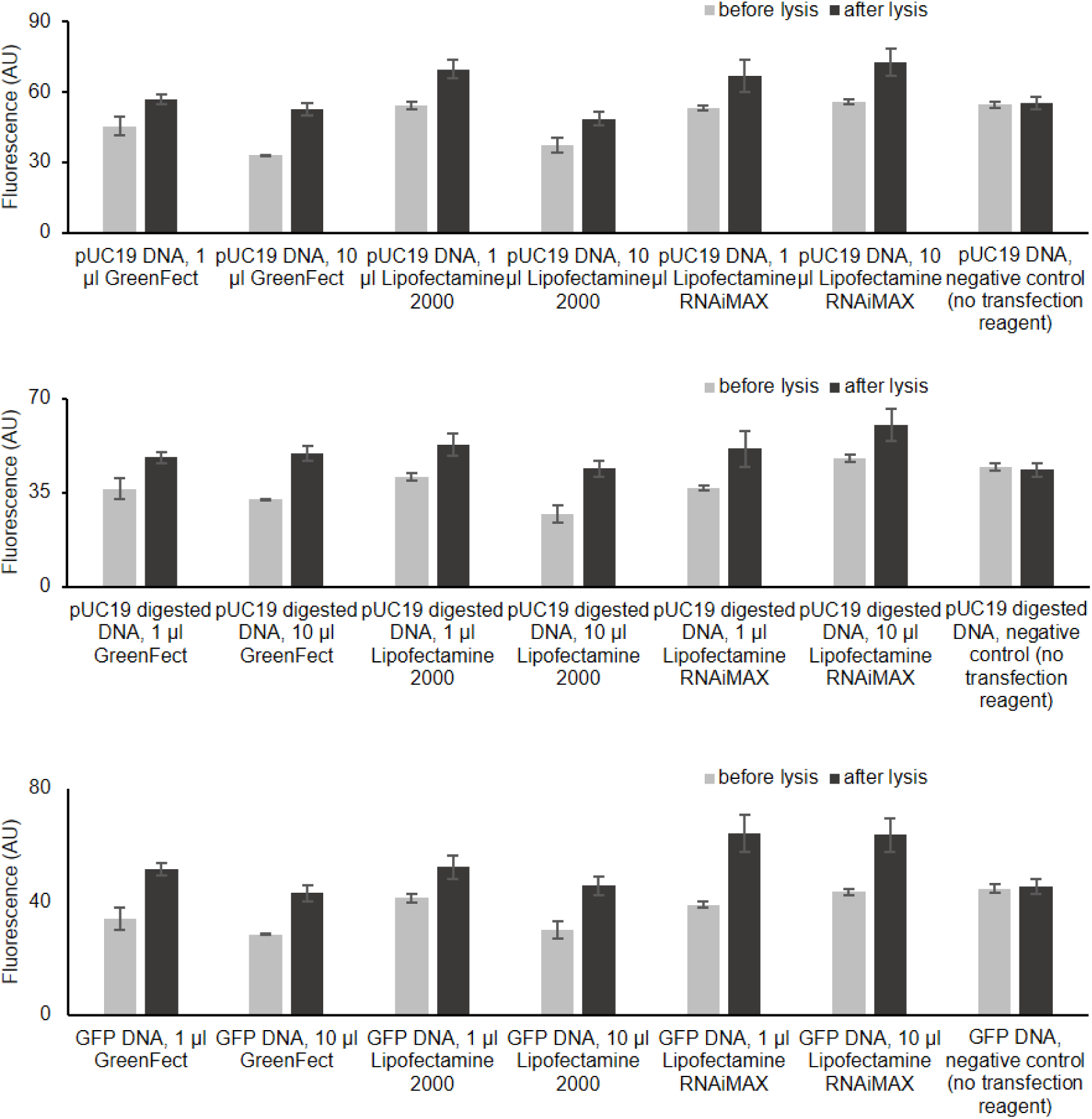
Transfection experiments using Diamond Dye lysis assay, three top performing transfection reagents, DNA payload.

**Supplementary Figure 36.**
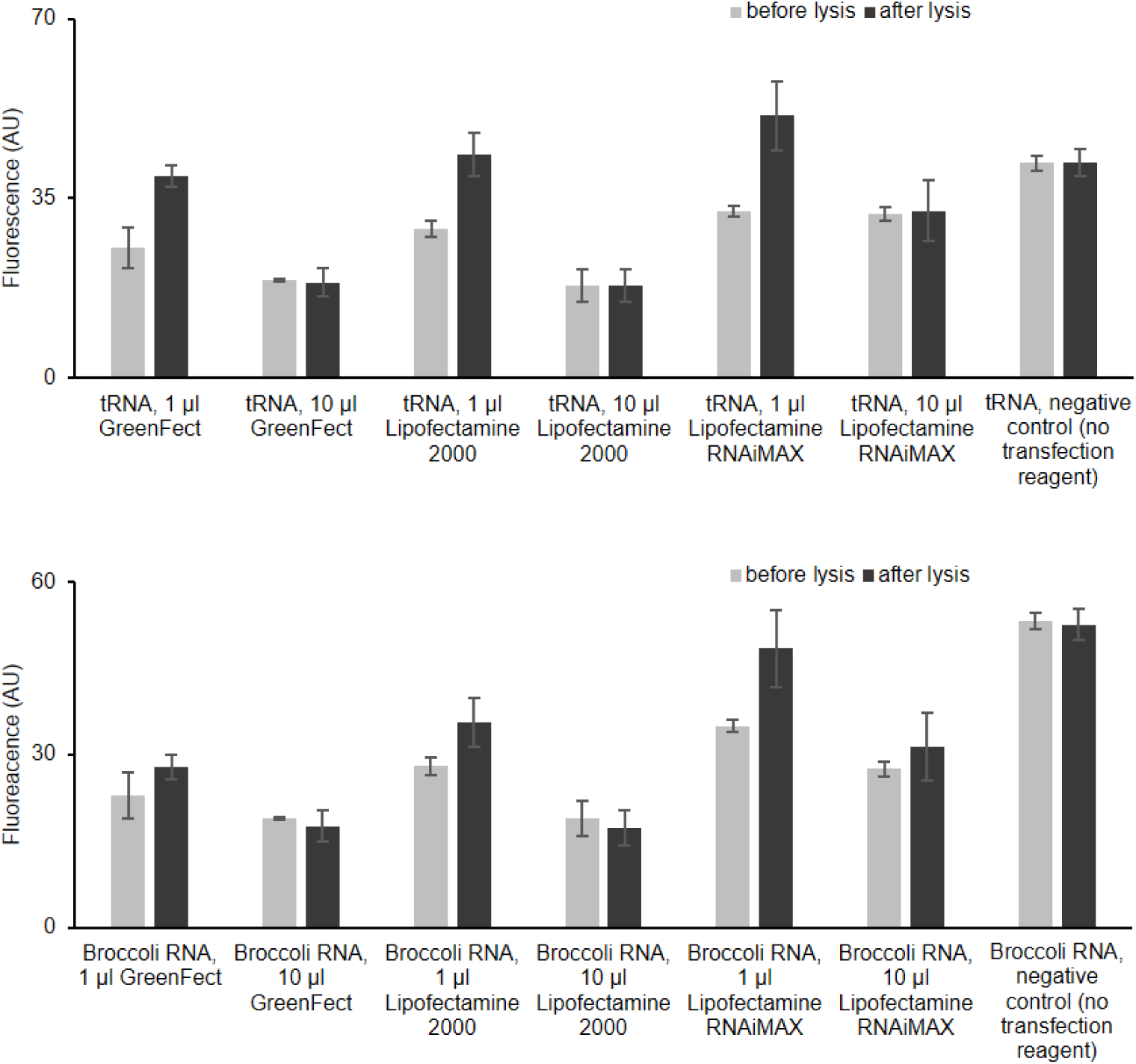
Transfection experiments using Diamond Dye lysis assay, three top performing transfection reagents, RNA payload.

## Tables

Please submit tables at the end of your text document (in Word or TeX/LaTeX, as appropriate). Tables that include statistical analysis of data should describe their standards of error analysis and ranges in a table legend.

**Supplementary Table 1.**
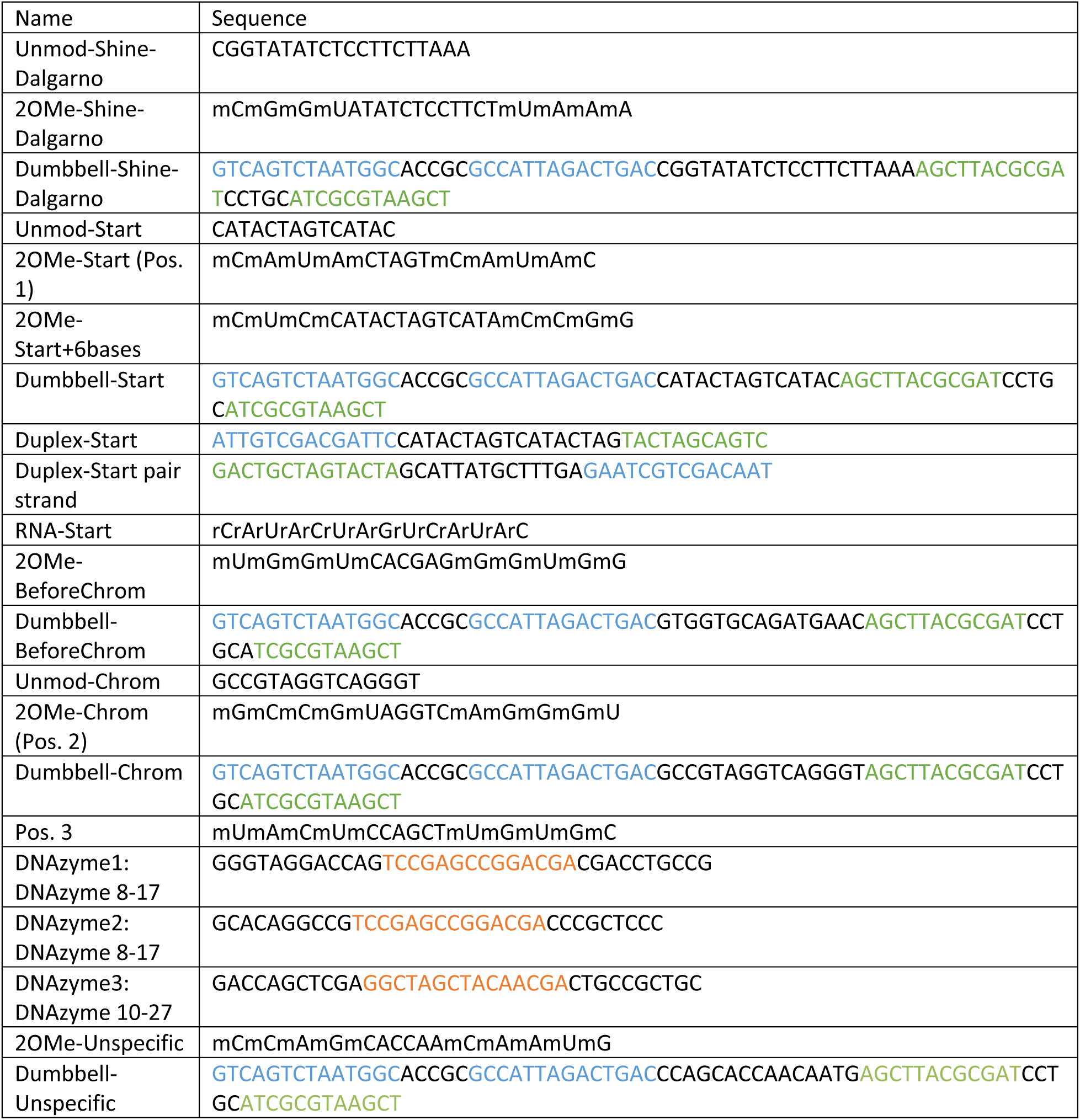
Silencing oligo sequences. • Dumbbell oligo: Same color sequences (green and blue) hybridizes to form dumbbell structure. • Duplex-ends oligo: Oligo 8 and 9 hybridize to form circle structure, the same color sequences (green and blue) are complemental sequences. Oligo 8 contains the complement sequence of deGFP mRNA. • DNAzyme: Orange sequences are the catalytic sequence for the DNAznyme. Black sequences are complementary to deGFP mRNA. • Oligo table

**Supplementary Table 2.** Initial transfection reagents test with POPC/cholesterol liposomes.

| Transfection reagent | Did it work?<br>(qualitative<br>assessment<br>only) | Gel analysis |
| --- | --- | --- |
| Mirus TransIT-LT1 | no | Supplementary Fig. 16,<br>Supplementary Fig. 19 |
| GenePORTER Gold Transfection Reagent | no | Supplementary Fig. 17,<br>Supplementary Fig. 18 |
| GenePORTER Gold with Diluent | no | Supplementary Fig. 17,<br>Supplementary Fig. 18 |
| GenePORTER Gold with Enhancer | no | Supplementary Fig. 17,<br>Supplementary Fig. 18 |
| GenePORTER 3000 Transfection Reagent | yes | Supplementary Fig. 18 |
| GenePORTER 3000 with Diluent | yes | Supplementary Fig. 18 |
| Canvax Canfast Transfection Reagent | no | Supplementary Fig. 20 |
| ICAFectin 441 (K) DNA Transfection<br>Reagent | no | Supplementary Fig. 20 |
| ICAFectin 441 (P) DNA Transfection<br>Reagent | no | Supplementary Fig. 20 |
| ICAFectin mRNA (K) mRNA Transfection<br>Reagent | yes | Supplementary Fig. 20 |
| ICAFectin mRNA (P) mRNA Transfection<br>Reagent | no | Supplementary Fig. 20,<br>Supplementary Fig. 21 |
| LiFect293 DNA Transfection Reagent | no | Supplementary Fig. 22 |
| GreenFect Transfection Reagent | yes | Supplementary Fig. 21,<br>Supplementary Fig. 22 |
| GreenFect with Enhancer Reagent | no | Supplementary Fig. 21,<br>Supplementary Fig. 22 |
| Avalanche-Omni Transfection Reagent | no | Supplementary Fig. 19 |
| Lipofectamine RNAiMAX Reagent | yes | Supplementary Fig. 16,<br>Supplementary Fig. 19 |
| Lentifectin Reagent | no | Supplementary Fig. 16 |
| DNAfectin Plus Transfection Reagent | no | Supplementary Fig. 20 |
| Mirus TransIT-mRNA | no | Supplementary Fig. 23 |
| Mirus mRNA Boost | no | Supplementary Fig. 23 |
| Mirus TransIT-X2 | no | Supplementary Fig. 23 |
| Lipofectamine 2000 Reagent | yes | Supplementary Fig. 23 |
| ViaFect Transfection Reagent | no | Supplementary Fig. 21,<br>Supplementary Fig. 24 |

**Supplementary Table 3.**
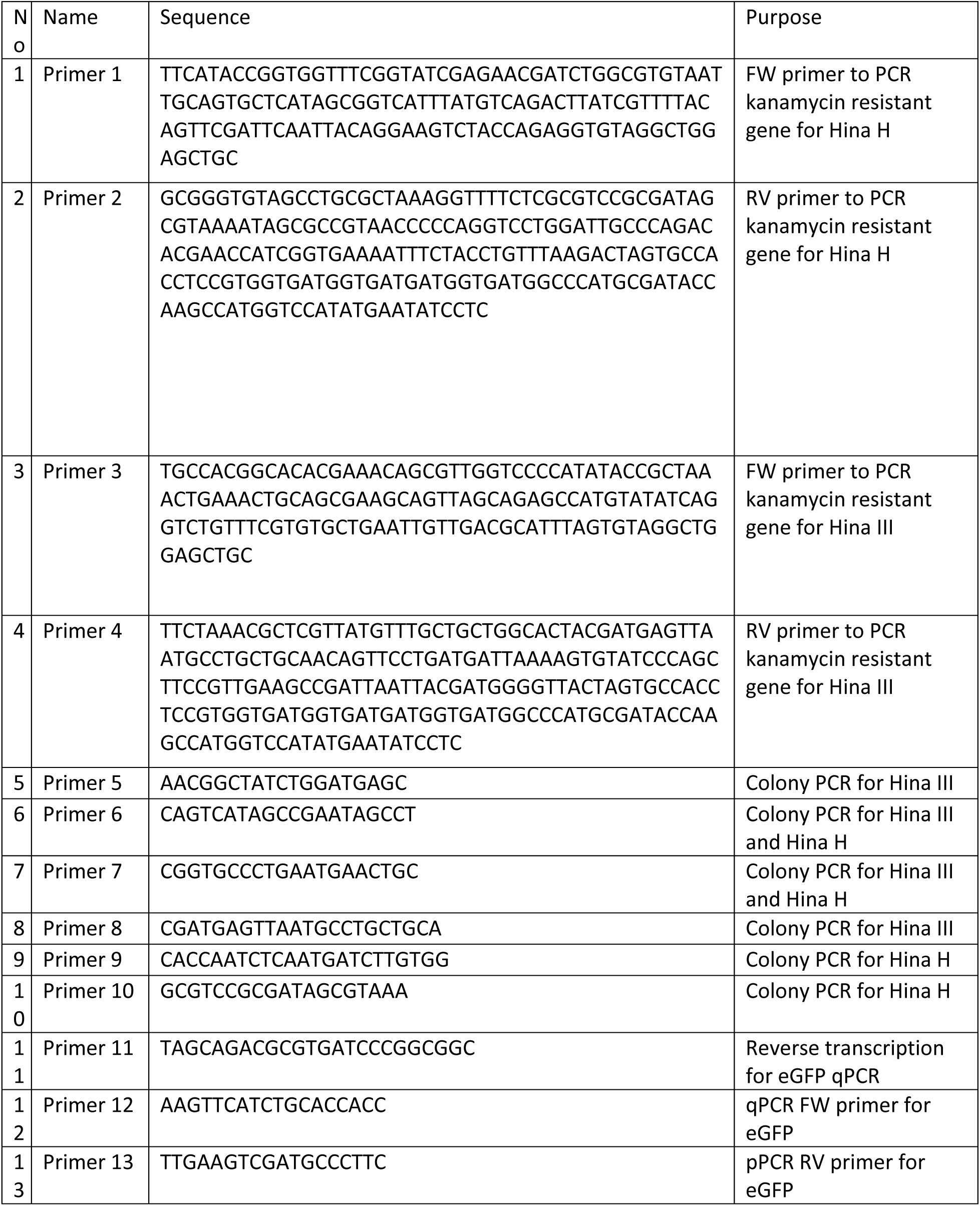
Primer sequences.

**Supplementary Table 4.**
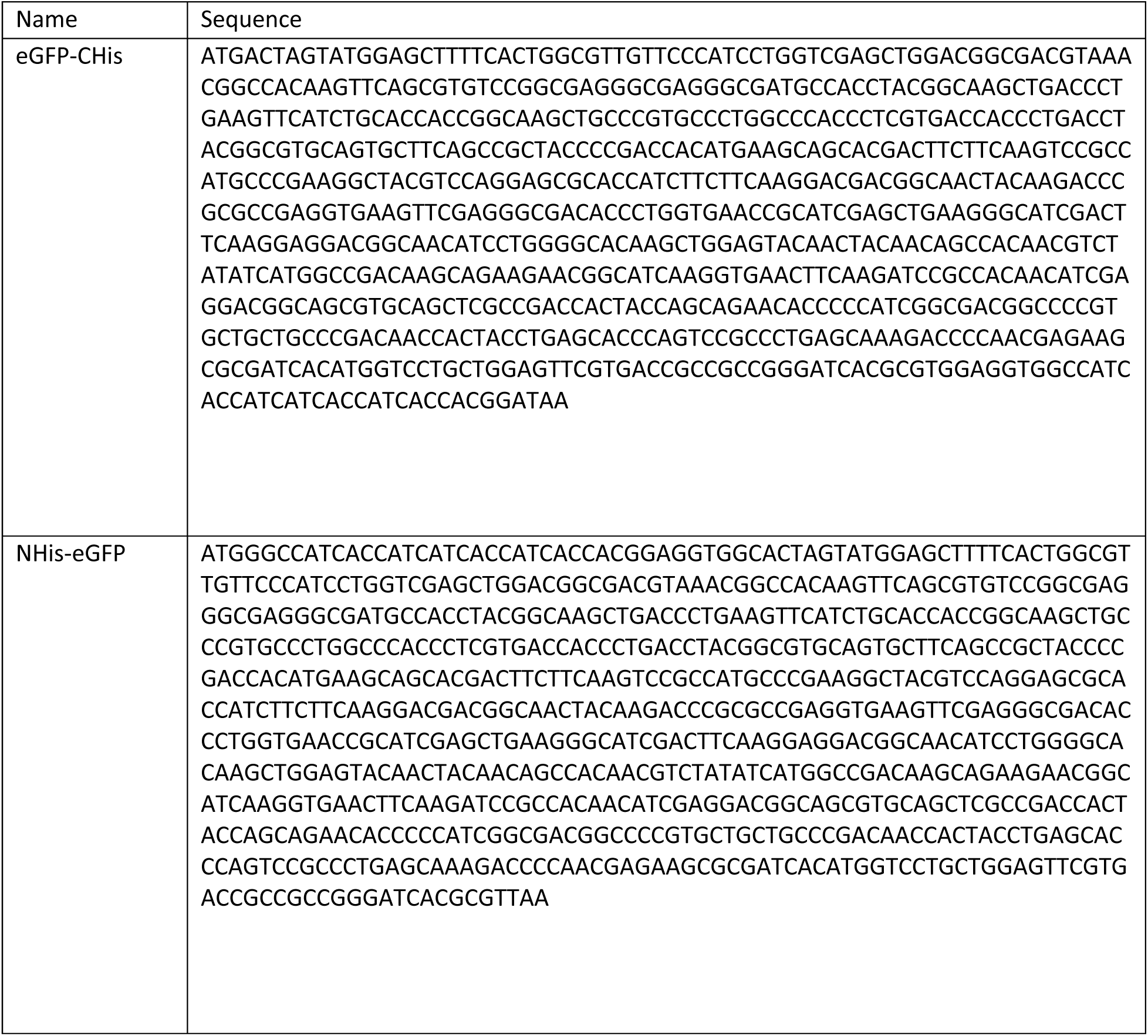
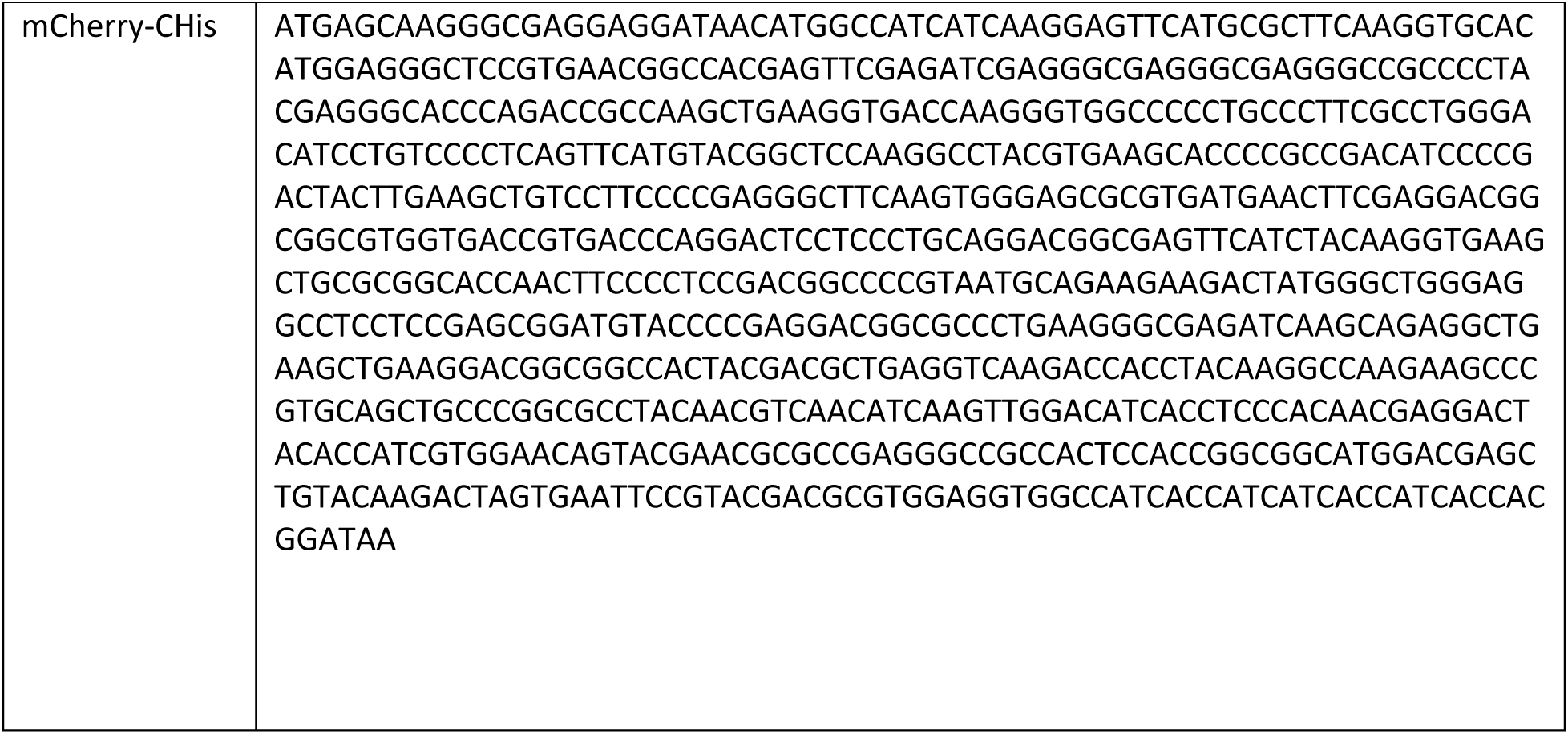
Gene sequences.

